# Rosindol: A fluorogen for the quantitative measurement of reactive oxygen species in living cells

**DOI:** 10.64898/2026.06.04.730197

**Authors:** Andrew Monnone, Molly Nicknish, Juan Jaramillo Montezco, Christine Sanganoo, Nalin Aggarwal, Arista Luong, Scott Schaus, Mark Grinstaff

## Abstract

Reactive oxygen species (ROS) are key mediators of disease, yet accurate characterization in living systems remains challenging because current probes lack oxidation specificity and produce nonlinear, pH-dependent signals. Here we introduce Rosindol, a novel thioacetal-based fluorogenic probe that overcomes these limitations. Rosindol undergoes an *umpolung* oxidation in the presence of ROS to generate fluorescence, displaying dose-linear responses to H_2_O_2_, O_2_•⁻, OH•, and HOCl with minimal background signal. Unlike conventional probes, Rosindol is pH-independent, photostable, water soluble, and agnostic to glucose concentration, esterase expression, and ambient oxygen. Validation in human cells—including PMA-stimulated neutrophils and SOD knockout models—confirms accurate detection of cytosolic and mitochondrial ROS. In pancreatic cancer cells, Rosindol reveals a fourfold increase in mitochondrial O_2_•⁻ generation capacity via Complex I of the electron transport chain. Glucose stimulation induces twofold higher ROS generation in malignant cells, highlighting a connection between Warburg metabolism and the etiology of oxidative stress in pancreatic cancer. These studies illustrate the utility of Rosindol to provide valuable insight to oxidative stress processes in complex biological environments.

## Main

The advent of artificial intelligence is transforming the analysis of complex biological data in research and medicine. However, the development of useful predictive algorithms relies upon datasets generated from accurate measurement tools. Consequently, the quality of tools used to measure biological processes is more relevant than ever. Molecular probes are measurement tools that provide precise characterization of disease processes in biomedical research and diagnostic medicine. Over the past century, the emergence of such probes has offered unprecedented spatiotemporal detection of pathological cellular analytes and metabolites^9,10^. Reactive oxygen species^5,6^ (ROS, Fig. 1a) including superoxide (O_2_•⁻) and hydrogen peroxide (H_2_O_2_) are endogenous metabolites upregulated in disease processes such as inflammation, ischemia, and malignancy^1–4^. While these small molecule oxidants mediate a wide span of physiologic processes on the nanomolar scale, significant ROS elevations induce a pathological shift in the cell environment termed oxidative stress^7^. Although the literature describing oxidative stress biology is abundant, precise characterization of the etiology, magnitude, and role of ROS in disease remains challenging due to the limitations of current ROS measurement probes^8^.

**Figure 1.**
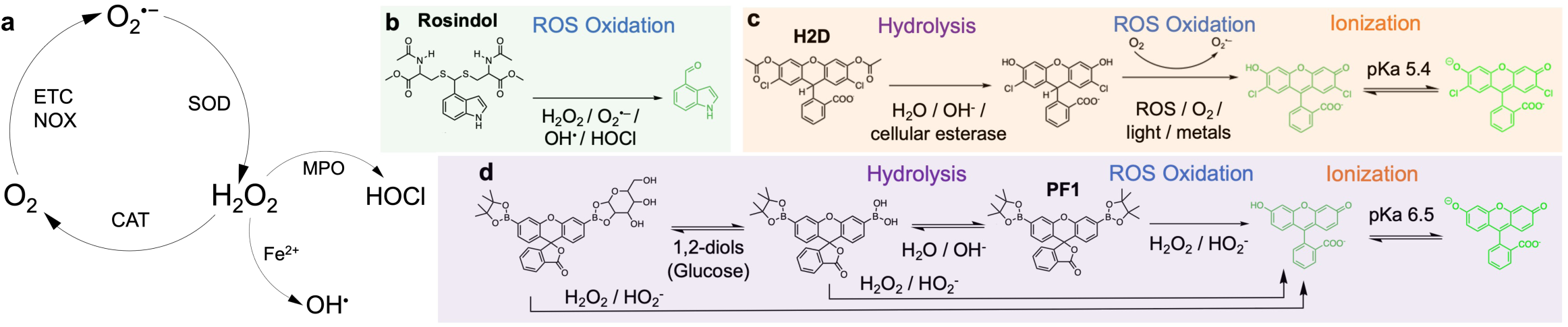
Reactive oxygen species (ROS) and chemical strategies for their measurement. **a.** Several key biological ROS analytes and their abbreviated metabolism pathways. **b.** Thioacetal probe Rosindol. **c.** Dihydro-xanthene ester probe 2’7’-dichlorodihydrofluorescein diacetate, “H2D”. **d.** Boronate-xanthene ester probe Peroxyfluor-1, “PF1”.

Introduced 60 years ago, dihydro-xanthene esters (e.g. 2’7’-dichlorodihydrofluorescein diacetate^11^, “H2D”, Fig. 1c) have remained the most widely used probes for measuring ROS *in vitro* despite lacking ROS-specific reactivity or signal intensity^12–14^. The inventors of H2D never proposed its use in human cells, and this practice has been formally discouraged since 1999 by the editors of *Free Radical Biology and Medicine*^15^. The activation of H2D *in vitro* depends on hydrolysis via cellular esterase, and its signal is therefore tied to expression levels of the enzyme^16,17^. This confounds measurement of ROS across different cell types, and in processes displaying abnormal esterase regulation such as malignancy^18,19^. Decades of studies and a 2022 consensus on ROS measurement have further established that H2D activation is not specific to ROS, showing sensitivity to pH, light, oxygen (O_2_), and metals (Murphy *et al.*)^8^. The fluorogenic trigger is a triphenyl network with broken conjugation, rendering the probe highly prone to autoxidation^11^ rather than selectively responsive to strong oxidants. Nonetheless, H2D does not react directly with the key ROS analyte H_2_O_2_, relying first on its conversion to OH• via peroxidases and/or ferrous iron^13^. Subsequent oxidation of the probe via OH• is self-amplifying, coupled to O_2_ reduction and the generation of O_2_•⁻. This results in nonlinear signal increases with changes in ROS concentration, impeding quantitative determinations^15^. Despite these significant limitations, H2D measurements underpin the vast majority of all conclusions on oxidative stress published. Its properties have distorted our perception since the early foundations of ROS biology theory, and its unabated use continues to warp the findings of emerging studies.

Boronate-xanthene esters (Chang et al^20^, e.g. Peroxyfluor-1, “PF1”, Fig. 1d) undergo hydrolytic deprotection directly via H_2_O_2_, eliminating the esterase activation step that confounds H2D measurements. However, the boronate / H_2_O_2_ reaction is nonoxidative, and minimally detects O_2_•⁻, OH•, and HOCl v. The mechanism is nucleophilic and pH-dependent^21^, facing competition from an H_2_O-driven pathway that yields a boronic acid and a 1,2-diol. This dynamic equilibrium^22–24^ results in a mix of species that react at different rates with H_2_O_2_, presenting several challenges for signal specificity *in vitro*. Once liberated, boronic acids readily complex with abundant biological 1,2-diols^25–27^ such as glucose, a well-documented propensity leveraged as a glucose-sensing strategy in cells^28–30^. Altered reactivity of the glucose-bound probe with H_2_O_2_ induces a signal intensity dependent on glucose concentration. Glucose is present at mM concentrations in all cells and its regulation varies considerably, as seen in the ‘Warburg’ metabolism observed in malignant cells^31–33^.

While H2D and PF1 use alternate mechanisms to detect ROS, both rely on xanthene fluorophores to report signal, which possess several measurement drawbacks. Xanthene fluorescence intensity is a function of fluorophore ionization, and thus exquisitely sensitive to changes in pH^34–37^. This results in ROS readouts that are influenced by variables such as subcellular location, cell culture/analysis practices, and the presence of metabolic pathology. For example, malignant cells display abnormalities in pH regulation due to the Warburg-driven accumulation of lactic acid^38,39^. Further, xanthenes exhibit low aqueous solubility and require use of the organic cosolvent dimethyl sulfoxide (DMSO), which induces ROS generation in cells^40–42^. Their narrow Stokes shift can lead to signal saturation, especially when using microplate readers^43–45^. Xanthene fluorophores are also sensitive to photobleaching, requiring dark handling and remaining vulnerable to irradiation by excitation wavelength photons during fluorescence measurement^46–49^. When used for reporting ROS *in vitro*, xanthene signals include significant noise from competing sources.

Herein, we describe Rosindol - a ROS-responsive fluorogen that exhibits signal specificity, pH independence, aqueous solubility, and broad species detection range. A key chemical design feature in Rosindol (Fig. 1b) is the thioacetal functional group, which is stable to a wide variety of chemical environments while cleaving readily in the presence of strong oxidants^50,51^. Rosindol undergoes an *umpolung* oxidation to generate the fluorophore indole-4-carboxaldehyde^52^ (IND4), displaying dose-linear fluorescence to concentrations of H_2_O_2_, O_2_•⁻, OH•, and HOCl, four key ROS analytes. Detection of ROS generation/inhibition controls in human cells validates Rosindol to accurately measure ROS *in vitro*. Assay of malignant and benign pancreatic cells using Rosindol reveals insights into ROS metabolism within a complex pathological environment featuring variations in pH regulation, O_2_ and glucose metabolism, and esterase expression. Rosindol demonstrates benchtop stability to light and air, quantitative measurements in cells, and outperforms xanthene probes, warranting application in biological systems.

## Results and Discussion

### Design and fluorogenic mechanism

From a performance perspective, we envisioned a fluorogen that demonstrates: 1) time and concentration dependent response to specific ROS (H_2_O_2_, O_2_•⁻) in the μM range or less; 2) signal independent of pH, O_2_, glucose, or esterase activity; 3) a wide Stokes shift and low background signal; 4) biocompatibility [≥ 95% cell viability after 24 hours at assay concentration]; 5) sufficient aqueous solubility to avoid the necessity of DMSO; 6) stability to ambient light, air, and hydrolysis; and, 7) compatibility with a broad range of detection modalities including microplate reader, confocal microscope, and flow cytometer. These criteria align with the principles outlined by the ROS measurement consensus of Murphy *et al.*^8^

From a design perspective, we identified thioacetal oxidation as an underutilized mechanism for ROS-specific probe activation. Classically employed as an aldehyde protecting group in organic synthesis^53^, here we leverage the thioacetal as an oxidation-dependent fluorogenic trigger. Thioacetal protection of a fluorescent aldehyde imposes an *umpolung* polarity inversion^54,55^ that attenuates donor-acceptor intramolecular charge transfer^56,57^, thereby extinguishing fluorescence. Oxidation of the thioacetal via ROS regenerates the aldehyde, reversing the polarity gradient and restoring fluorescence. Critically, mechanistic characterization of this oxidation indicates the sulfur byproduct is a disulfide rather than a radical-scavenging thiol^51^ (Fig. S33). We synthesized a series of thioacetal probes based upon this *‘ROS umpolung’* principle to examine structure-activity relationships. Optimization of the aldehyde led to the green-emitting IND4 as an ideal fluorophore for three reasons. First, the compound demonstrates no measurable cytotoxicity at concentrations up to 100 μM for 24 hours (Fig. S21). Second, the fluorophore undergoes no ionization across the physiological pH range, displaying pH-independent signal intensity (Fig. 2g, top panel). Third, it possesses a wide Stokes shift, allowing precise detection of emission wavelength light without interference from excitation wavelength light (Fig. 2b). Rosindol is the thioacetal of IND4 and the thiol O-Me-N-acetyl-L-cysteine. We synthesized Rosindol via N-protection of IND4 as its 9-fluorenylmethyl carbamate followed by addition of the thiol under electrophilic conditions (Fig. S18, 48% yield over three steps). Relative to IND4, Rosindol displays a hypsochromic shift in its absorbance peak to the UV region, and the complete absence of an emission peak (Fig. 2b).

**Figure 2.**
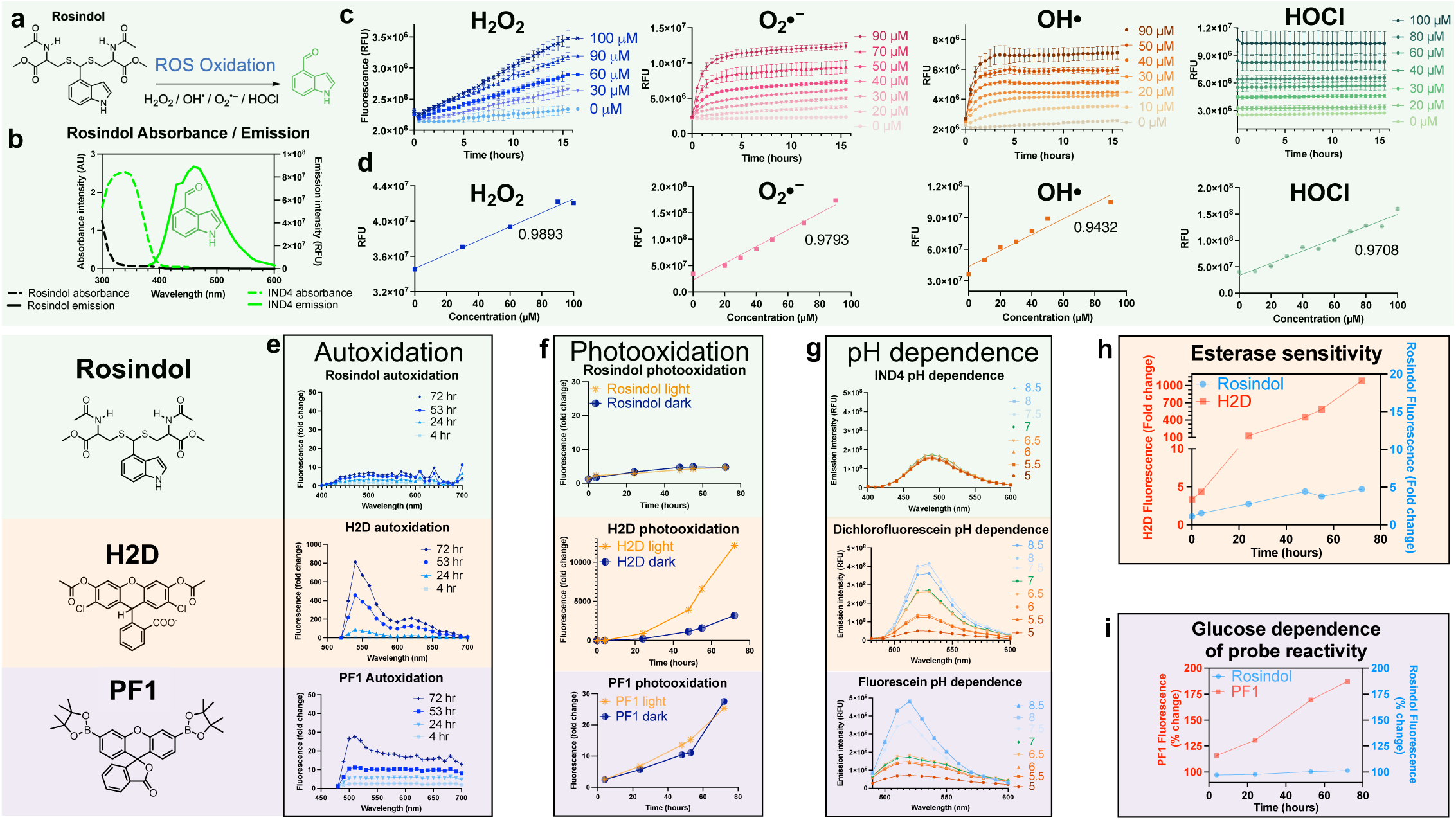
Oxidation and signal specificity of Rosindol. **a.** Rosindol oxidation reaction. **b.** Absorbance/emission spectra of 10 μM Rosindol and IND4. **c.** Time-dependent fluorescence of 100 μM Rosindol in the presence of 0 – 100 μM ROS analytes measured via microplate reader. **d.** Linear regression analysis of Rosindol fluorescence and ROS concentration. **e.** Time-dependent emission spectrum intensity of ROS probes in solutions maintained in the dark and exposed to air measured via microplate reader. **f.** Time-dependent emission peak wavelength intensity of ROS probes exposed to ambient laboratory lighting. **g.** Emission spectrum intensity of 10 μM solutions of probe fluorophores over the pH range 5 – 8.5. **h.** Time-dependent emission peak wavelength intensity of Rosindol and H2D in the presence of 23 units of porcine liver esterase. **i.** Time-dependent emission peak wavelength intensity of Rosindol and PF1 in the presence of 20 mM glucose and 100 μM H_2_O_2_.

### Oxidation via ROS and signal specificity

We next characterized the concentration-dependent dynamic range and temporal response of Rosindol to H_2_O_2_, O_2_•⁻, OH•, and HOCl, four key biological ROS analytes. Fluorescence of 100 μM aqueous Rosindol was measured across concentrations of 0 - 100 μM ROS over 16 hours via microplate reader (Fig. 2c). Both the rate of signal increase and signal intensity are proportional to the oxidative potential of the measured ROS analytes. While fluorescence plateaus in the presence of OH• (+2.3 V)^13^ after three hours, reaction with H_2_O_2_ (+0.32 V) is still progressing at 15 hours. After calculating the area under each curve (AUC) and performing linear regression analysis, a dose-linear signal is observed for all four species (Fig. 2d), indicating two merits for Rosindol as a probe. First, linear interpolation allows calibration of *in vitro* signals against known concentrations of ROS, enabling quantitative measurements in the presence of adequate positive controls. This ability stands in contrast to the self-amplifying oxidation of the xanthene probe H2D, which generates a nonlinear signal precluding straightforward quantification. Second, reactivity with four important ROS analytes constitutes a wider detection range than either xanthene probe, capturing a greater percentage of total cellular ROS during measurement.

To examine signal specificity, we characterized the ROS-independent activation of Rosindol and the commercially available xanthene ROS probes H2D and PF1. We measured the emission spectrum intensity of the probes over 72 hours in the presence of ambient O_2_ (autoxidation), light (photooxidation), porcine liver esterase, and β-D-glucose, as well as the emission intensity of the probe fluorophores across a range of pH values. 100 μM Rosindol demonstrates significantly lower rates of autoxidation compared to 10 μM solutions of either xanthene probe (Fig. 2e) and does not display H2D’s susceptibility to photooxidation (Fig. 2f), illustrating the robust specificity of thioacetal oxidation as a mechanism for ROS measurement. The presence of esterase alone is sufficient to increase H2D fluorescence by several orders of magnitude in a time-dependent manner, highlighting the low oxidation specificity of this probe (Fig. 2h). In contrast, Rosindol displays no increase in fluorescence in the presence of esterase. H_2_O_2_ detection via boronate probe PF1 demonstrates significantly higher signal in the presence of 20 mM glucose (DMEM cell culture media = 25 mM glucose), while Rosindol signal shows no such dependence (Fig. 2i, S34, S35). While xanthene fluorophore ionization couples emission intensity to pH, IND4 signal displays minimal variation across the pH range of 5 – 8.5 (Fig. 2g). Altogether, these studies illustrate the advantages of Rosindol for the specific measurement of ROS in the presence of significant competing oxidative and biological variables.

### Validation for the measurement of ROS in human cells

According to the guiding principles of the 2022 ROS measurement consensus of Murphy *et al.*^8^, a small molecule probe should be validated to detect specifically generated ROS, negative controls, knockdowns, inhibitors, and radical scavengers, and do so in a plausible manner regarding time and dose. Using this framework, we manipulated ROS in human cells and characterized the ability of Rosindol to detect the perturbations. We first assessed the cytotoxicity of Rosindol to ensure the probe itself would not induce ROS through cellular injury/death pathways^58,59^. After 24 hours of exposure, the IC_50_ of Rosindol is ≥1 mM via MTS assay in human pancreatic and kidney cells (Fig. 3a). No cytotoxicity is observed at 100 μM, allowing live cell measurements at this concentration for up to 24 hours.

**Figure 3.**
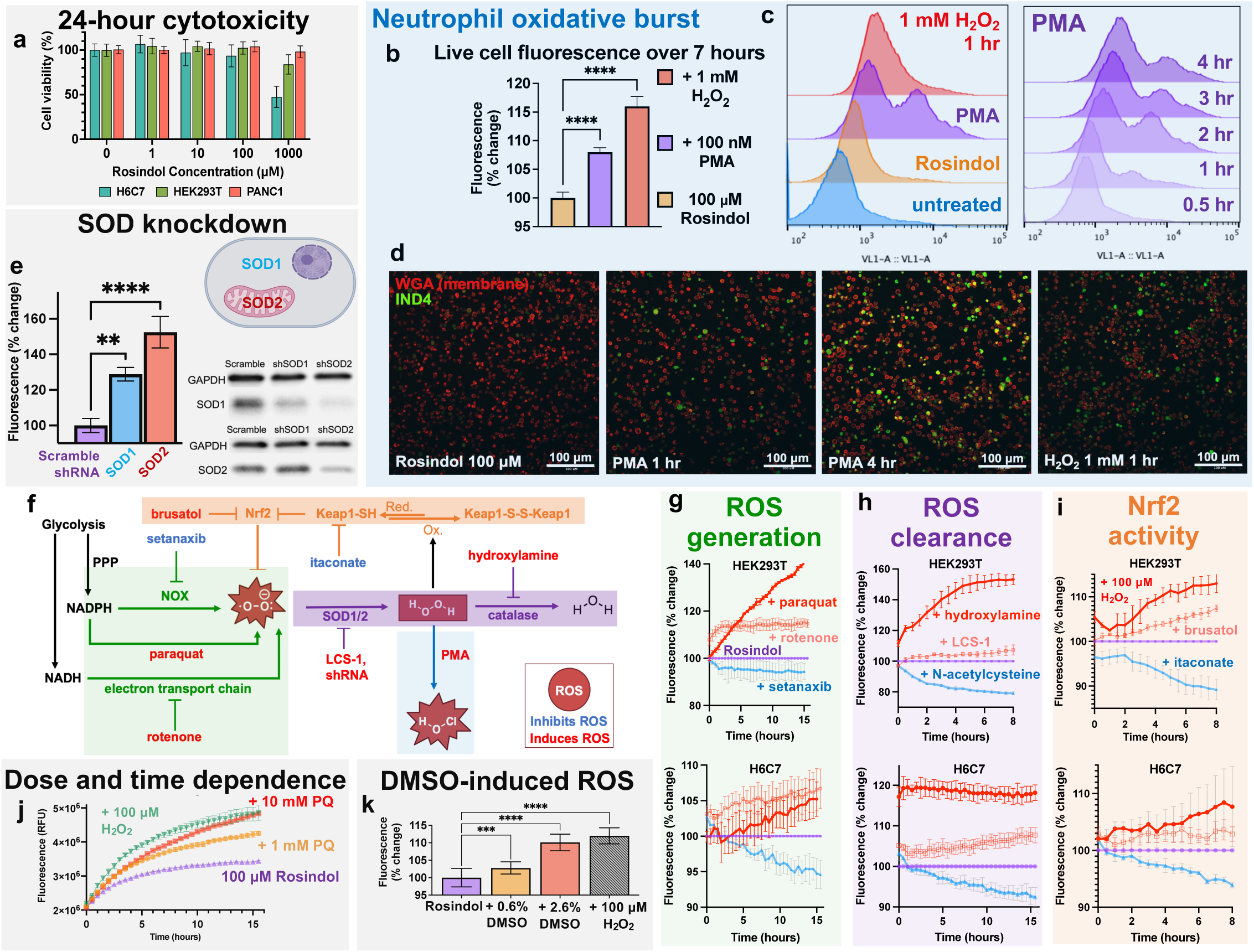
Validation for the measurement of ROS in human cells. **a.** 24-hour cytotoxicity of Rosindol measured via MTS cell viability assay in 3 mammalian cell lines. **b.** Rosindol signal of PMA-stimulated neutrophils over 7 hours via microplate reader. **c.** Rosindol signal of PMA-stimulated neutrophils via flow cytometer over 4 hours. **d.** Rosindol signal of PMA-stimulated neutrophils via confocal microscope over 4 hours. **e.** Rosindol signal (via flow cytometry at 1 hour) and SOD expression (via Western blot) of HEK293T cells following SOD1 and SOD2 shRNA knockdown. **f.** Canonical pathways of cellular ROS metabolism examined in human cells. **g.** Rosindol fluorescence measurements in HEK293T (top panel) and H6C7 cells (bottom panel) via microplate reader following treatment with small molecule manipulators of O₂•⁻ generation pathways. **h.** Fluorescence measurements via microplate reader following inhibition of O_2_•⁻ and H_2_O_2_ clearance pathways. **i.** Fluorescence measurements via microplate reader following inhibition of Nrf2 and its repressor Keap1. **j.** Time and dose dependence of Rosindol signal in the presence of 1 and 10 mM paraquat (PQ) in HEK293T cells via microplate reader. **k.** Dose dependence of Rosindol signal in the presence of DMSO in HEK293T cells via microplate reader. Significance was determined via ordinary one-way ANOVA using a normal (Gaussian) distribution assumption and multiple comparisons. P values were calculated and graphed using the following designations: P ≤ 0.0001 = **** ; P ≤ 0.001 = *** ; P ≤ 0.01 = ** ; P ≤ 0.05 = * ; P ≥ 0.05 = NS (not significant), by Tukey’s post-hoc test.

Among the panoply of differentiated human cells, neutrophils provide the quintessential example of robust biological ROS generation. In response to picomolar-level stimulation from pathogenic metabolites, neutrophils undergo “oxidative burst” to generate bactericidal quantities of O_2_•⁻, H_2_O_2_, and HOCl^56^. After differentiating human promyeloblast (HL-60) cells into neutrophils^60–62^, we treated the cells with 100 μM Rosindol and 100 nM of the potent stimulatory molecule phorbol 12-myristate 13-acetate (PMA)^63^. We included neutrophils treated with 1 mM H_2_O_2_ and Rosindol were as a positive control. Rosindol fluorescence significantly increases over time in the PMA-treated neutrophils via microplate reader, confocal microscope, and flow cytometer (Fig. 3b-d). Live cell kinetic fluorescence indicates the signal in PMA-activated neutrophils is equivalent to approximately 500 μM H_2_O_2_ over seven hours (Fig. 3b). Flow cytometry demonstrates a bimodal distribution of fluorescing cells, revealing a brightly emitting secondary population that peaks two hours after PMA treatment (Fig. 3c). These studies indicate the probe accurately detects the biological ROS generation of human cells *in vitro*, with potential utility as a diagnostic neutrophil stain for identifying bacterial infection^9,64^.

The majority of biological ROS are generated in mitochondria^65–67^, and thus the ability to measure mitochondrial ROS is essential for the accuracy and utility of an *in vitro* probe^68,69^. However, small molecules can face transport barriers that prevent efficient uptake in the highly regulated and selective mitochondrial compartments^70,71^. Thus, we assessed the ability of Rosindol to measure mitochondrial ROS via translational repression of the cytosolic and mitochondrial isoforms of the antioxidant enzyme superoxide dismutase^72^ (SOD1/2). We transfected HEK293T cells with plasmids encoding SOD1, SOD2, or non-targeting scrambled shRNA, and measured fluorescence via flow cytometry 1 hour after Rosindol treatment. We confirmed SOD knockdown via Western blot for SOD1/2. Compared to the scrambled shRNA control, ROS signal increases in cells transfected with either SOD1 and SOD2 plasmids (Fig. 3e), indicating Rosindol measures both cytosolic and mitochondrial sources of ROS. A larger increase in signal is observed in cells transfected with the SOD2 plasmid, which likely reflect the increased generation of ROS in the mitochondria relative to the cytosol, and/or the result of a combination SOD1/SOD2 knockdown facilitated by SOD2 shRNA, as indicated by Western blot.

We next assessed the ability of Rosindol to measure perturbations in canonical pathways known to govern cellular ROS metabolism (Fig. 3f). Using commercially available small molecule inhibitors, we manipulated ROS metabolism pathways in human kidney (HEK293T) and pancreatic (H6C7) cells and measured the change in Rosindol signal via live cell kinetic fluorescence. We induced O₂•⁻ generation via rotenone inhibition of Complex 1 of the mitochondrial electron transport chain (ETC)^73,74^, Complex 1 redox cycling via paraquat^75,76^, and inhibition of NADPH oxidase (NOX)^77–80^ (Fig. 3g). We inhibited O_2_•⁻ and H_2_O_2_ clearance via inhibition of SOD1 and catalase^81–83^ (Fig. 3h), and manipulated antioxidant regulation via inhibition of nuclear factor erythroid 2-related factor 2 (Nrf2)^84–86^ and its repressor Kelch-like ECH-associated protein 1 (Keap1)^87,88^ (Fig. 3i). We treated cells with 100 μM H_2_O_2_ and 1 mM N-acetylcysteine^89,90^ as direct positive and negative ROS controls (Fig. 3h, 3i). The results show Rosindol detects increases/decreases in ROS that would be expected following each manipulation of the ROS metabolism pathways (Fig. 3g-i) in a dose and time dependent manner (Fig. 3j). Compared to benign pancreatic cells (Fig. 3g-i, bottom panels), fully transformed HEK293T cells (Fig. 3g-i, top panels) show increased ROS signals and more exaggerated responses to pathway manipulations. These studies illustrate that Rosindol accurately detects changes in cellular ROS following manipulation of ROS metabolism pathways *in vitro*.

The organic cosolvent DMSO is required for the use of xanthene probes and shows a dose-dependent effect on ROS signal (Fig. 3k), emphasizing the benefit of aqueous solubility as a probe quality. The aqueous solubility of Rosindol is approximately 1 mM (Fig. S19, S22), allowing *in vitro* measurements at 100 μM without the necessity of DMSO.

### ROS measurements in malignant and benign pancreatic cells

Oxidative stress affords several traits that support the survival and proliferation of malignant cells^91,92^ and there is substantial reported evidence of elevated ROS in cancer^93,94^. Pancreatic ductal adenocarcinoma (PDAC) is the deadliest common malignancy in the United States, associated with elevated ROS that contribute to aberrant signaling, genomic instability, and unrestrained proliferation^95–97^. However, the etiology of elevated ROS in PDAC remains obscured. The complex malignant cell environment features anomalies in pH, O_2_^98^, glucose metabolism, and esterase expression, overwhelming the limited performance of conventional ROS probes. In contrast, Rosindol demonstrates independence to these variables (Fig. 2e-i), and accurately detects ROS metabolism manipulations *in vitro* (Fig. 3g-i). With these properties established, we used Rosindol to measure ROS levels in malignant and benign pancreatic cells and characterize distinctions in malignant ROS metabolism.

Compared to benign pancreatic cells, Rosindol shows significantly higher signal in malignant cells via microplate reader (Fig. 4a), flow cytometer (Fig. 4b), and confocal microscope (Fig. 4c), affirming the presence of elevated ROS in PDAC. Live cell kinetic fluorescence indicates ROS levels over 10 hours are 85% and 40% higher in malignant Panc1 and BxPC-3 cells than those in benign H6C7 cells, respectively (Fig. 4a). We then examined the ability of Rosindol to selectively identify malignant cells in a malignant/benign co-culture via their increased ROS levels. Live cell imaging indicates significantly increased Rosindol signal in malignant Panc1 cells (Fig. 4d, red stain) compared to benign dermal fibroblasts (Fig. 4d, blue stain). We repeated the experiment using the xanthene probe H2D, which conversely demonstrates significantly higher activation in the fibroblasts and does not identify the malignant cells in the co-culture (Fig. 4e, S29, S30).

**Figure 4.**
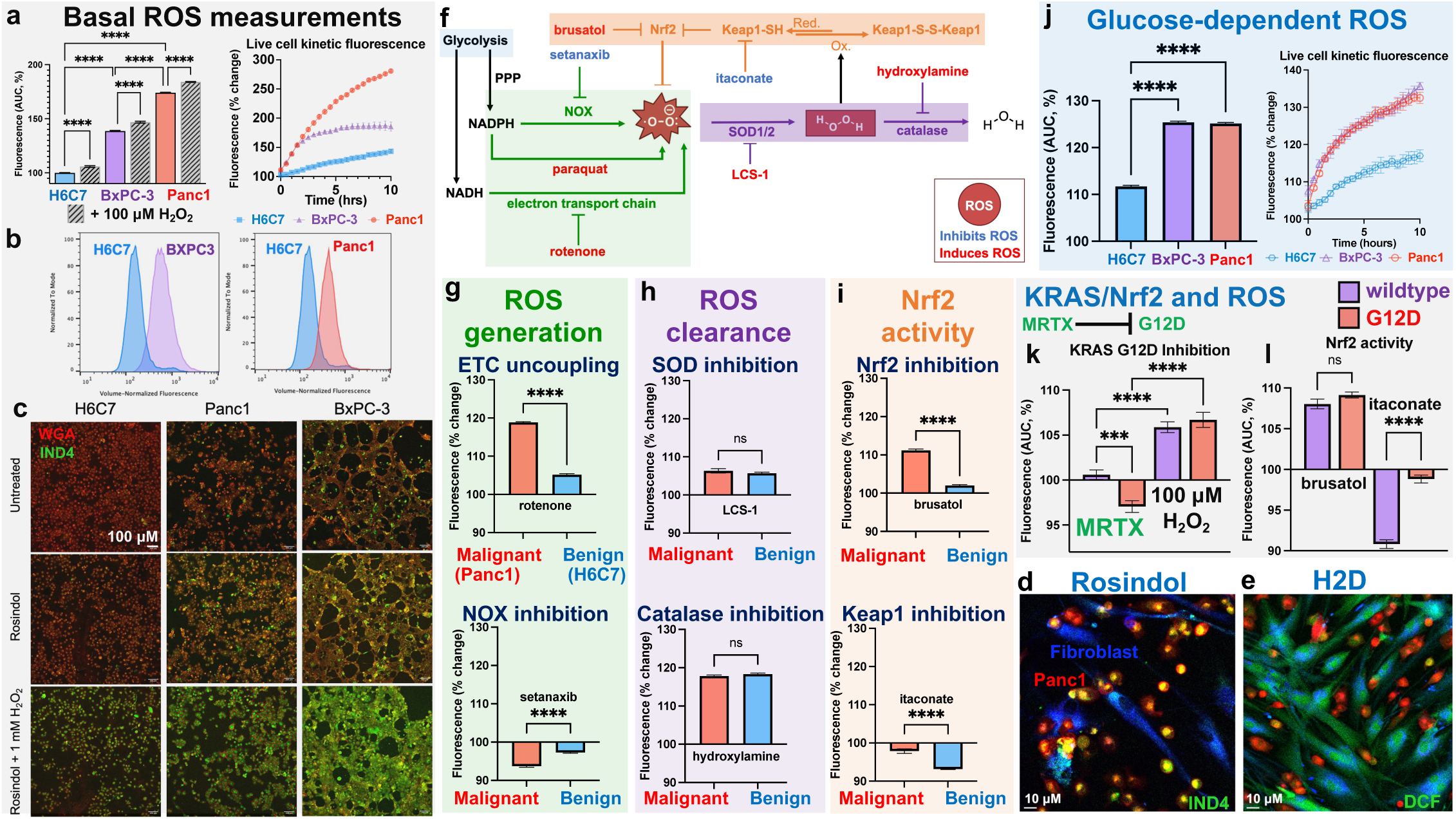
Measurement of ROS in malignant and benign pancreatic cells. **a.** ROS measurements of malignant and benign pancreatic cells over 10 hours via microplate reader. **b.** ROS measurements via flow cytometry 1 hour after Rosindol treatment. **c.** ROS measurements via live cell confocal microscopy four hours after Rosindol treatment. **d.** Microscopy of a Panc1/human dermal fibroblast co-culture stained with Rosindol. **e.** Microscopy of a Panc1/human dermal fibroblast co-culture stained with H2D. **f.** Canonical pathways of ROS metabolism examined in malignant and benign pancreatic cells. **g.** Manipulation of O_2_•⁻ generation pathways in Panc1 and H6C7 cells. **h.** Inhibition of O_2_•⁻ and H_2_O_2_ clearance pathways in Panc1 and H6C7 cells. **i.** Inhibition and induction of Nrf2 activity in Panc1 and H6C7 cells. **j.** Glucose-dependent ROS generation in malignant and benign cells. **k.** ROS generation in Panc1 (KRAS^G12D^) and BxPC-3 (KRAS^WT^) following treatment with 100 nM of the G12D inhibitor MRTX1133 for nine hours. **l.** ROS generation following Nrf2 inhibition and induction in Panc1 and BxPC-3 cells. Significance was determined via ordinary one-way ANOVA using a normal (Gaussian) distribution assumption and multiple comparisons. P values were calculated and graphed using the following designations: P ≤ 0.0001 = **** ; P ≤ 0.001 = *** ; P ≤ 0.01 = ** ; P ≤ 0.05 = * ; P ≥ 0.05 = NS (not significant), by Tukey’s post-hoc test.

Next, we manipulated canonical pathways of ROS metabolism in Panc1 and H6C7 cells to identify distinctions in malignant ROS regulation (Fig. 4f). No differences are seen in O_2_•⁻ / H_2_O_2_ clearance via SOD / catalase (Fig. 4h), suggesting these pathways are not responsible for PDAC oxidative stress. Conversely, malignant cells show a greater disruption in ROS generation following NOX inhibition (Fig. 4g, bottom panel), suggesting higher basal NOX activity. Nrf2 antioxidant activity also appears higher at baseline in malignant cells, along with a diminished capacity for further induction via Keap1 inhibition (Fig. 4i). These findings are consistent with a compensatory upregulation of antioxidant pathways in response to oxidative stress, which has previously been observed in malignant cells^99–101^. Most notably, malignant cells exhibit a four-fold higher capacity to generate mitochondrial O_2_•⁻ following inhibition of Complex 1 of the electron transport chain (ETC, Fig. 4g, top panel).

### Glucose metabolism and oxidative stress in PDAC

The precursors of mitochondrial O_2_•⁻ are O_2_ and nicotinamide adenine dinucleotide hydride (NADH)^102–104^. Malignant cells forgo the consumption of O_2_ for ATP generation, relying on elevated levels of glucose and NADH for energy production (the ‘Warburg Effect’^31–33^). The definitive impact of Warburg metabolism on ROS levels is complex and debated, with rational arguments supporting both pro-oxidative and ROS-balancing roles^105,106^. We hypothesized that due to the increased provision of available NADH and O_2,_ malignant cell glucose metabolism might facilitate elevated O_2_•⁻ generation. To examine this hypothesis, we stimulated malignant and benign cells with 200 mM glucose and measured the change in Rosindol signal via live cell kinetic fluorescence. Following glucose treatment, ROS generation increases by a two-fold higher magnitude in malignant Panc1 and BxPC-3 cells than benign H6C7 cells (Fig. 4j). ROS generation in malignant cells is higher following glucose stimulation than any other pathway manipulation examined, including inhibition of ETC Complex 1 (Fig. 4g, top panel) and catalase (Fig. 4h, bottom panel). These results indicate that glucose is a potent stimulus for ROS generation in malignant cells, highlighting a connection between Warburg metabolism and the etiology of oxidative stress in PDAC cells.

### K-ras mutation and oxidative stress in PDAC

A mutation in the K-ras oncogene (KRAS)^107,108^ is present in the majority of pancreatic cancers, and its conclusive role in oxidative stress remains unresolved. Several studies suggest KRAS mutation is associated with higher ROS levels in cancer cells^109–112^. Conversely, a 2022 study by Fan *et al.* reports that KRAS^G12D^ mutation eliminates ROS in PDAC by driving upregulation of Nrf2^113^. However, the authors did not measure lower ROS levels in KRAS^G12D^ cells. Using the xanthene probe H2D without a positive control, they observed no differences in ROS between KRAS^G12D^ and wildtype (WT) cells (Fig. 1E^113^). Further, the authors proposed no biochemical mechanism of interaction to connect the G12D protein to Nrf2 upregulation, which has previously been characterized as a compensatory response to increased ROS levels^99–101^, rather than a preemptive driver of lower basal ROS levels. In our own studies, significantly higher ROS generation occurs in KRAS^G12D^ cells (Panc1) than KRAS^WT^ cells (BxPC-3) via live cell kinetic fluorescence (Fig. 4a), suggesting that the KRAS^G12D^ mutation correlates to higher ROS levels. This prompted our curiosity to re-examine the relationship between KRAS^G12D^ mutation, Nrf2 activity, and ROS in the presence of inhibitors and positive controls.

We treated KRAS^G12D^ and KRAS^WT^ cells with the G12D protein inhibitor MRTX1133^114,115^ for 9 hours, followed by replacement of the media and live cell kinetic fluorescence measurements over 10 hours via Rosindol. Compared to uninhibited cells, ROS generation decreases in KRAS^G12D^ cells alone (Fig. 4k, ‘MRTX’, Fig. S27), suggesting the function of the G12D protein is associated with higher ROS levels. Nrf2 activity was examined in both cell lines by measuring ROS generation following inhibition and induction (Fig. 4l). KRAS^G12D^ cells show significantly less response to Nrf2 induction via itaconate, suggesting increased activity at baseline relative to KRAS^WT^ cells. Nrf2 inhibition via brusatol results in equal levels of ROS generation in both cell lines, suggesting Nrf2 activity alone is unlikely to explain differences in their basal ROS levels.

Based on these results, we propose that KRAS^G12D^ mutation in PDAC cells is associated with higher ROS levels and upregulated Nrf2 activity, with the latter representing a compensatory response to the former. It is likely the prior findings of Fan *et al.* are a consequence of the low signal specificity of H2D or premature autoxidation of the probe, which cannot be ruled out in the absence of a positive control in their ROS measurement. We encourage further exploration of the role of KRAS^G12D^ mutation in oxidative stress to fully elucidate its function in PDAC pathology and progression. Such studies will provide valuable insight to inform the emerging clinical use of G12D inhibitors as a treatment modality for PDAC^116,117^.

### Concluding Remarks

Since their discovery in the latter half of the 20^th^ century, ROS have increasingly been recognized as significant mediators of biological disease processes. Attaining the full benefit of this insight requires improving our understanding of ROS, and hence our measurement tools. Attention to this need is already being noted, with recent innovations including the development of genetically engineered proteins that provide unprecedented sensitivity and spatial resolution to detect H_2_O_2_ in cells^118^. However, utilizing such systems requires altering the genetic expression of the target cell line via plasmids or viral vectors prior to ROS measurement, which itself induces oxidative stress in cells^119,120^ and poses experimental challenges as far as assay timing, cost, and convenience.

As a small molecule, Rosindol is ready for immediate use in unmodified cell lines and is compatible with a wide range of fluorescence measurement modalities. Its low cytotoxicity allows for the observation of oxidative stress processes in living cells over 24 hours, and its dose-linear response to key ROS analytes facilitates straightforward quantitative measurements. The signal is agnostic to relevant oxidative and biological variables including pH, O_2_, light, glucose concentration, and esterase activity, and it accurately detects biological ROS generation and inhibition controls in human cells. While it is unlikely for any single probe to be appropriate for all measurement tasks, we advocate for thioacetal oxidation as a valuable chemical strategy for the measurement of reactive oxygen species *in vitro.* We next intend to further our investigation into ROS-responsive thioacetals via the development of probes optimized for the measurement of ROS *in vivo*.

## Methods/Experimental Section

### Synthetic materials and characterization methods

All chemicals and reagents were purchased from commercial suppliers and used without further purification. NMR spectra were recorded on a 500 MHz Varian NMR spectrometer. UPLC mass spectra were recorded on a Waters Acquity UPLC with evaporative light scattering detector (ELSD), SQ mass spectrometer, and Waters 2996 PDA (photodiode array) detector. High-resolution mass spectra were recorded on a Waters Xevo G3 QTof.

### Synthesis of (9*H*-fluoren-9-yl)methyl 4-formyl-1*H*-indole-1-carboxylate (Fig. S1)

To an oven-dried 100mL round bottom flask, 4-indole-carboxaldehyde (1g, 6.89mmol, 1eq) and 30mL of anhydrous methylene chloride were added under nitrogen atmosphere. The solution was stirred at 0℃ via ice bath, followed by the addition of sodium hydride (824mg, 34.4mmol, 5eq). The mixture was stirred for 30 minutes, followed by the addition of fluorenylmethyloxycarbonyl chloride (3.55g, 13.79mmol, 2eq) and warming to 40℃. The reaction was monitored by TLC in a solution of cyclohexane: ethyl acetate (3:1) to confirm conversion of starting material. After 24 hours, the reaction was cooled to room temperature, quenched with 10 mL of deionized water, and the organic layer was collected. The aqueous layer was extracted 3x with 10 mL methylene chloride. The combined organic extracts were washed with brine, dried with magnesium sulfate, and concentrated under vacuum to yield an amber oil. The crude mixture was purified via flash column chromatography on silica gel (eluent: 18:82 ethyl acetate: cyclohexane) and the product was obtained as a light pink crystalline solid (2.02g, 5.51mmol, 80% yield).^1^H NMR (d_6_-DMSO, 500 MHz): δ 10.18 (s, 1H), 7.93 (d, *J* = 7.5 Hz, 2H), 7.79 – 7.71 (m, 4H), 7.55 (s, 1H), 7.43 (t, *J* = 7.4 Hz, 2H), 7.35 (td, *J* = 7.4, 1.2 Hz, 2H), 7.29 – 7.22 (m, 2H), 5.02 (d, *J* = 4.9 Hz, 2H), 4.54 (t, *J* = 5.0 Hz, 1H). ^13^C NMR (d_6_-DMSO, 500 MHz): δ 193.46, 150.53, 143.90, 141.47, 135.25, 129.33, 128.89, 128.61, 128.52, 128.32, 127.80, 125.32, 124.59, 120.71, 120.68, 107.28, 69.03, 46.87. UPLC-MS: calculated for [M+H^+^]: 368.4, found: 368.326.

### Synthesis of (9*H*-fluoren-9-yl)methyl 4-(4-acetamido-10-(methoxycarbonyl)-3,12-dioxo-2-oxa-6,8-dithia-11-azatridecan-7-yl)-1*H*-indole-1-carboxylate (Fig. S6)

An oven dried 50mL round bottom flask was charged with (9*H*-fluoren-9-yl)methyl 4-formyl-1*H*-indole-1-carboxylate (500mg, 1.36mmol, 1eq), N-acetyl-L-cysteine methyl ester (0.529mg, 2.99mmol, 2.2eq), and 15mL of anhydrous methylene chloride under nitrogen atmosphere. The solution was stirred at 0℃ via ice bath, followed by the dropwise addition of boron trifluoride diethyl etherate (0.629mL, 5.45mmol, 4eq), and the solution developed an orange color. The reaction was warmed to room temperature and monitored by TLC (1:1, cyclohexane: ethyl acetate). After 3 hours the reaction was quenched with 5 mL of a saturated aqueous solution of sodium bicarbonate, and the organic layer was collected. The aqueous layer was extracted 3x with 10 mL methylene chloride. The combined organic extracts were washed with brine, dried with magnesium sulfate, and concentrated under vacuum to yield a crude oil. The crude mixture was purified via flash column chromatography on silica gel (eluent: 90:10 ethyl acetate: cyclohexane) and the product was obtained as a white crystalline solid (797mg, 1.13mmol, 83% yield). ^1^H NMR (d_6_-DMSO, 500 MHz): δ 8.32 (dd, *J* = 17.4, 8.3 Hz, 2H), 7.94 (d, *J* = 7.6 Hz, 2H), 7.75 (d, *J* = 7.5 Hz, 2H), 7.58 (d, *J* = 3.7 Hz, 1H), 7.44 (t, *J* = 7.5 Hz, 2H), 7.35 (t, *J* = 7.4 Hz, 2H), 7.25 (d, *J* = 7.5 Hz, 2H), 7.06 (t, *J* = 8.0 Hz, 1H), 6.94 (d, *J* = 3.8 Hz, 1H), 5.57 (s, 1H), 4.99 (d, *J* = 5.1 Hz, 2H), 4.53 (t, *J* = 5.0 Hz, 1H), 4.44 (q, *J* = 7.4, 6.8 Hz, 2H), 3.60 (s, 3H), 3.55 (s, 3H), 3.01 (dd, *J* = 13.5, 5.5 Hz, 1H), 2.82 – 2.76 (m, 2H), 2.73 (dd, *J* = 13.4, 8.9 Hz, 1H), 1.82 (s, 3H), 1.79 (s, 3H). ^13^C NMR (d_6_-DMSO, 500 MHz) δ 171.56, 171.47, 169.84, 169.76, 150.68, 143.97, 141.48, 134.98, 132.11, 128.62, 128.30, 127.78, 125.91, 125.31, 124.66, 122.06, 120.70, 114.81, 107.29, 68.76, 52.47, 52.25, 52.00, 50.89, 46.89, 33.82, 33.63, 22.66. HR-MS: calculated for [M+Na^+^]: 726.79, found: 726.1951.

### Synthesis of methyl *S*-(((2-acetamido-3-methoxy-3-oxopropyl)thio)(1*H*-indol-4-yl)methyl)-*N*-acetylcysteinate (Rosindol, Fig. S12)

To an oven dried 25mL round bottom flask, (9*H*-fluoren-9-yl)methyl 4-((4*R*,10*R*)-4-acetamido-10-(methoxycarbonyl)-3,12-dioxo-2-oxa-6,8-dithia-11-azatridecan-7-yl)-1*H*-indole-1-carboxylate (600mg, 0.85mmol, 1eq) and 10mL anhydrous dimethylformamide (DMF) were added under nitrogen atmosphere, followed by 2mL piperidine. The reaction was stirred at room temperature whilst being monitored via TLC (1:1, cyclohexane: ethyl acetate). After 3 hours, the reaction mixture was evaporated under reduced pressure, and the crude oil was redissolved in 50 mL methylene chloride, washed sequentially with 5 mL deionized water and brine, and concentrated under vacuum. The crude product was purified via flash column chromatography on silica gel (eluent: 100% ethyl acetate) to yield a white crystalline solid (298mg, 0.62mmol, 73%).

^1^H NMR (d_6_-DMSO, 500 MHz): δ 11.22 – 11.16 (m, 1H), 8.35 (dd, *J* = 22.7, 7.9 Hz, 2H), 7.38 – 7.31 (m, 2H), 7.11 – 7.02 (m, 2H), 6.65 (t, *J* = 2.5 Hz, 1H), 5.54 (s, 1H), 4.53 – 4.44 (m, 2H), 3.62 (s, 3H), 3.58 (s, 3H), 3.02 (dd, *J* = 13.5, 5.7 Hz, 1H), 2.82 (d, *J* = 7.0 Hz, 2H), 2.75 (dd, *J* = 13.4, 8.9 Hz, 1H), 1.84 (s, 3H), 1.81 (s, 3H).^13^C NMR (d_6_-DMSO, 500 MHz): δ 171.68, 171.58, 169.86, 169.77, 136.68, 130.54, 126.15, 125.60, 121.15, 118.06, 111.93, 100.53, 52.46, 52.31, 52.14, 52.03, 33.86, 33.64, 22.69, 22.66. HR-MS: calculated for [M+H^+^]: 482.6, found: 482.1414.

### Aqueous solubility (Fig. S19, S20)

A solubility standard curve was prepared by dissolving Rosindol in DMSO at 4 different concentrations from 0.5 – 5 mg/mL and analyzing the samples via UPLC. Evaporative light-scattering detector (ELSD) peak area was then plotted vs. sample concentration in mg/mL to yield a calibration curve. Linear regression analysis indicates a linear relationship between ELSD peak area and sample concentrations over the range of 0.5 – 5 mg/mL (R^2^ = 0.9856). Aqueous solubility was then assessed by adding 5 mg of the tested compound to 1 mL of UltraPure distilled water (Invitrogen) in a 5 mL glass scintillation vial, vortexing the mixture for 30 seconds, and allowing the solution to equilibrate for 24 hours. The sample was then filtered via 0.4 μM PTFE syringe filter, and ELSD peak area was analyzed via UPLC. The DMSO peak area standard curve was then used to calibrate the aqueous sample peak area and determine the aqueous solubility of Rosindol (Fig. S19). This process was repeated using H2D to determine its comparative aqueous solubility (Fig. S20). For practical aliquoting and experimentation, microliter quantities of DMSO were used to prepare concentrated stocks of Rosindol, which were then diluted in water to working concentrations of ≤ 0.04% DMSO for fluorescence assays. Solutions of 0.04% DMSO demonstrated the same results in cells as fully aqueous Rosindol (Fig. S22).

### Measurement of photophysical properties (Fig. 2b)

10 μM aqueous solutions of compounds were prepared using UltraPure distilled water. 100 μL aliquots of each solution were then added to 96-well black-walled plates for analysis, and the absorbance and emission spectra were recorded using a Spectramax iD3 spectrophotometer and Softmax Pro.

### ROS-responsive kinetic fluorescence assay (Fig. 2c, 2d)

100 μM aqueous solutions of Rosindol were prepared using UltraPure distilled water, and 100 μL aliquots were distributed into a 96-well black-walled plate. 10 – 100 μmol of each ROS analyte was then added to wells in triplicates, and a clear adhesive plastic film was fixed to the top of the plate to minimize evaporative loss from the wells. The fluorescence intensity (excitation 360 nm, emission 490 nm) was then measured every 30 minutes for 16 hours using a Spectramax iD3 spectrophotometer set in kinetic mode, with low gain, read from the top face of the plate. ROS standards were prepared as follows: H_2_O_2_ and HOCl were purchased as 30% and 8% aqueous solutions respectively and diluted using UltraPure distilled water as necessary. OH• was generated *in situ* via addition of 1 equivalent of Iron (II) Bromide (FeBr_2_) to H_2_O_2_. O₂•⁻ was generated *in situ* from H_2_O_2_ and sodium hydroxide (NaOH) according to the protocol established by Stoin et al^121^. Following collection of the kinetic fluorescence measurements, the data was imported to Graphpad Prism and plotted as the mean and standard deviation of the 3 replicates at each timepoint (Fig. 2c). The kinetic data was then examined via area under the curve (AUC) analysis and linear regression to yield concentration-dependent fluorescence calibrations (Fig. 2d).

### Signal specificity studies. (Fig. 2e-2i)

Aqueous solutions of Rosindol (100 μM), H2D (10 μM), and PF1 (10 μM) were prepared and 100 μL aliquots of each were distributed into a 96-well black-walled plate. The wells were covered with an adhesive plastic film to prevent evaporation and maintained 1 meter below an overhead fluorescent white light (CFL bulb) for 72 hours. A subset of the wells were tightly covered with an opaque foil to prevent exposure to the light. To assess autoxidation, the fluorescent emission intensity spectra were recorded at 4, 24, 53, and 72 hours via Spectramax iD3 spectrophotometer (Fig. 2e). To assess photooxidation, the emission intensity at peak spectral wavelength from the foil-covered and light-exposed wells were compared (Fig. 2f). To assess pH dependence of the probe fluorophores, 10 μM aqueous solutions of indole-4-carboxaldehyde (IND4), dichlorofluorescein (DCF), and fluorescein were prepared using UltraPure distilled water, and the emission intensity spectra were recorded. pH adjustments were then made using 1 M HCl and 1 M K_2_CO_3_ to achieve solutions of pH 5 - 8.5, followed by measurement of the fluorescent emission intensity spectra (Fig. 2g). The role of esterase on H2D and Rosindol activation was determined via the addition of 23 units of porcine liver esterase to 100 μM PBS solutions of the fluorogens and subsequent measurement of the emission intensity spectra over a period of 72 hours. The fluorescence intensity at the peak wavelength was then plotted versus time (Fig. 2h). The role of glucose on PF1 and Rosindol fluorescence was determined via measurement of the emission intensity spectra in the presence of 100 μM H_2_O_2_ with and without 20 mM β-D-glucose over a period of 72 hours. The fluorescence intensity at peak wavelength of the glucose-treated solution was then normalized to the glucose-free solution at each timepoint to examine glucose dependence (100% = no glucose dependence, fig. 2i).

### Mammalian cell culture

HEK293T and Panc1 cells were cultured in Dulbecco’s Modified Eagle Medium (DMEM) supplemented with 10% fetal bovine serum (FBS) and 1% penicillin/streptomycin (P/S). H6C7 cells were cultured in Keratinocyte serum-free media (KSFM) supplemented with bovine pituitary extract and human epidermal growth factor and 1% P/S. BxPC-3 cells were cultured in Roswell Park Memorial Institute (RPMI) medium supplemented with 10% FBS and 1% P/S. HL-60 cells were cultured in Iscove’s Modified Dulbecco’s Medium (IMDM) supplemented with 20% FBS and 1% P/S. Human dermal fibroblasts were cultured in Fibroblast Growth Medium (FGM). All cell cultures were maintained in T75 flasks at 37 C in 5% CO_2_ incubators, passaged at 80-100% confluence, and tested negative routinely for mycoplasma infection.

### Cell viability assays (Fig. 3a, S21)

Cells were seeded at 10,000 cells/well in black-walled cell culture-treated 96-well plates. After 24 hours, media was removed and replaced with 1 - 1000 μM of Rosindol in media in a series of tenfold dilutions. A subset of untreated wells were included to provide a 100% viable control. After 24 hours of exposure to Rosindol, media was removed and replaced with 100 μL of MTS reagent in cell media (1:5 dilution). After 1 hour, the absorbance at 490 nm was measured, and measurements were normalized to the untreated cells to gauge cell viability.

### Live cell kinetic fluorescence assay (general protocol)

Cells were seeded at 10,000 cells/well in a black-walled 96-well tissue culture plate. After 24 hours, media was removed and replaced with media containing 100 µM Rosindol. A subset of wells were replaced with Rosindol-free media to provide an untreated control. A second subset of wells were treated with 100 µM Rosindol and 100 µM H_2_O_2_ to provide a positive control for ROS. An adhesive clear plastic film was then fixed to the top of the wells to prevent evaporation. IND4 fluorescence intensity (excitation 360 nm, emission 490 nm) was then measured every 30 minutes for up to 16 hours using a Spectramax iD3 spectrophotometer maintained at 37 C, and the data was recorded using Softmax Pro 7. The Softmax Pro PMT gain was set to low and the measurements were recorded from the top of the wells. Each condition was assessed via 3 - 6 replicates per timepoint. After collection, data was imported to Graphpad Prism and plotted as the mean and standard deviation of the replicates at each timepoint (e.g. Fig. 3g, S24, S25). Kinetic data was then plotted as area under the curve (AUC) (e.g. Fig. 3b, S26) and standard error of the mean. Significance was determined via ordinary one-way ANOVA using a normal (Gaussian) distribution assumption and multiple comparisons. P values were calculated and graphed using the following designations: P ≤ 0.0001 = **** ; P ≤ 0.001 = *** ; P ≤ 0.01 = ** ; P ≤ 0.05 = * ; P ≥ 0.05 = NS (not significant), by Tukey’s post-hoc test.

### Confocal microscopy fluorescence (general protocol)

Cells were seeded at 10,000 cells/well in a 96-well black-walled glass-bottom tissue culture plate. After 24 hours, cells were treated with 100 µM Rosindol for 1 - 4 hours followed by treatment with 5 µg/mL wheat germ agglutinin (WGA)-Alexafluor647 as a cell membrane stain. A subset of wells were treated with 100 µM Rosindol and 1 mM H_2_O_2_ to provide a positive control for ROS. Cells were then fixed via 4% paraformaldehyde (PFA) or imaged live using a temperature (37° C) and CO_2_ (5%) regulated chamber. Cells were analyzed using an Olympus FV3000 microscope using 405 nm excitation to detect IND4 and 640nm excitation to detect WGA. For example PMT settings, see Supplemental Information Fig. S28.

### Flow cytometry fluorescence (general protocol)

Cells were seeded at 50,000 cells/well in a 24-well tissue culture plate. After 24 hours, cells were trypsinized and treated with 100 µM Rosindol for 1 - 4 hours. A subset of wells were treated with 100 µM Rosindol and 1 mM H_2_O_2_ to provide a positive control for ROS. At desired time points, aliquots of cells were removed and placed on ice, optionally fixed via 4% paraformaldehyde (PFA), and analyzed via an Attune NxT2 flow cytometer. Cells were analyzed via forward and side scatter gating and IND4 was detected using the VL1 laser (405 nm excitation). For example gating analysis, see Supplemental Information Fig. S31, S32.

### Measurement of ROS in human neutrophils (Fig. 3b-3d)

HL-60 cells were supplemented with 1% DMSO for 5 days to induce neutrophil differentiation^60–62^. Cells were then harvested and plated for fluorescence studies using microplate reader, confocal microscope, and flow cytometer as per the general protocols outlined above. Cells were treated with 100 µM Rosindol followed by 100 nM phorbol 12-myristate 13-acetate (PMA)^63^, and the fluorescent signal was measured over 1-4 hours via confocal microscope and flow cytometer, and over 7 hours via microplate reader. Cells treated with 100 µM Rosindol followed by 1 mM H_2_O_2_ were used as a positive control. Cells were fixed via 4% PFA prior to analysis via confocal microscopy and flow cytometry.

### RNA knockdown of SOD1/SOD2 (Fig. 3e)

HEK293T cells were seeded at 175,000 cells/well in a 12-well tissue culture plate pre-treated with poly-L-lysine. The cells were transfected the next day at 70-80% confluency. In each condition, cells were transfected using polyethylenimine (PEI)-plasmid complexes. To create the PEI-DNA complexes, PEI (0.323 g/L, Polysciences, 23966-2) were first combined with 0.15 M NaCl (4 parts PEI:21 parts NaCl). Next, 1000 ng DNA composed of (a) equal parts of three shRNA-encoding plasmids targeting three locations within the SOD1 transcript (Santa Cruz Biotechnology, sc-36523-SH), (b) equal parts of three shRNA-encoding plasmids targeting three locations within the SOD2 transcript (Santa Cruz Biotechnology, sc-41655-SH), (c) a degenerate shRNA-encoding plasmid which will not lead to the specific degradation of any cellular mRNA (Santa Cruz Biotechnology, sc-108060), or (d) an mCherry-encoding plasmid, were combined with 0.15 M NaCl and UltraPure distilled water to obtain 0.09 M NaCl and 20 ng/μL DNA. The two mixtures were then combined at a 1:1 ratio and allowed to form the PEI-DNA complex for 20 minutes at room temperature. Lastly, 100 μL of transfection reagent was dispensed per well (1000 ng). Forty-eight and seventy-two hours after transfection, HEK cells were harvested and treated with 100 µM Rosindol. Cells were then analyzed via flow cytometry 1 hour following treatment. Cell pellets were saved to perform western blot analysis.

### SOD1/2 Western blot (Fig. 3e)

Following shRNA knockdown and collection of cell pellets, HEK293T cells were lysed using 100 µL of Pierce RIPA buffer containing Halt™ Protease Inhibitor Cocktail in 1.5 mL Eppendorf microcentrifuge tubes that were kept under agitation for 30 minutes at 4°C. The samples were then spun in a microcentrifuge at 4°C for 20 minutes at 10,000 g and the supernatant was collected. Samples were quantified with the Pierce BCA Protein Assay Kit. 5 ug of protein per sample in 4x Laemelli Sample Buffer #1610747 were resolved on a Mini Protean TGX 4-20% gel at 100 volts for 80 minutes. Protein samples were transferred to a PVDF membrane at 100 volts for 90 minutes in chilled Towbin buffer. Membranes were blocked in 5% non-fat dry milk in Tris-Buffered Saline + 0.1% Tween20 (TBS-T) for 1 hour at room temperature. Membranes were incubated with the respective primary antibodies, diluted 1:1000 in blocking buffer, overnight at 4°C under constant agitation. The primary antibodies were rabbit-derived (GAPDH, #2118 from Cell Signaling Technology®, SOD1, #37385 from Cell Signaling Technology®, and SOD2, 835194RR from Proteintech®). To remove excess primary antibodies, the membranes were washed with TBS-T for 10 minutes 4 times. The membranes were then incubated with goat anti-rabbit HRP secondary antibodies, diluted 1:2000 in blocking buffer, for 1 hour at room temperature under constant agitation. To remove excess secondary antibodies, the membranes were washed with TBS-T for 10 minutes 4 times. The membranes were visualized using a ChemiDoc Imaging System and SuperSignal™ West Femto Maximum Sensitivity Substrate.

### ROS metabolism inhibitor studies in HEK293T and H6C7 cells (Fig. 3f-3i)

Cells were seeded at 10,000 cells/well in a 96-well black-walled plate as per the microplate reader general protocol detailed above. After 24 hours, media was removed and replaced with media containing 100 µM Rosindol, followed by the ROS generation/inhibition control compounds and immediate measurement via microplate reader. Paraquat and 4-octyl itaconate were added 9 hours prior to the fluorogen (16 hours after initial plating). Inhibitors were used in the following amounts: rotenone 200 nM, paraquat 10 mM, setanaxib 10 µM, LCS-1 10 µM, hydroxylamine 30 µM, brusatol 10 µM, 4-octyl itaconate 75 µM, N-acetylcysteine 1 mM, H_2_O_2_ 100 µM. Each condition was assessed via 3 - 6 replicates per timepoint. After collection, data was imported to Graphpad Prism and plotted as the mean and standard deviation of the replicates at each timepoint (e.g. Fig. 3g, S24, S25). Fluorescence in the presence of each control compound was normalized to cells treated with Rosindol alone to examine impact on ROS levels (see Supplemental Information Fig. S24-S26).

### ROS measurements in malignant and benign pancreatic cells (Fig. 4a-4c)

Panc1, BxPC-3, and H6C7 cells were seeded at 10,000 cells/well in 96-well plates for confocal microscopy and microplate reader studies, and 50,000 cells/well in a 24-well plate for flow cytometry, as per the general protocols outlined above. Live cell kinetic fluorescence was measured via microplate reader. Each condition was assessed via 3 - 6 replicates per timepoint. After collection, data was imported to Graphpad Prism and plotted as the mean and standard deviation of the replicates at each timepoint (e.g. Fig. 4a right-hand panel, S24, S25). Kinetic data was then plotted as area under the curve (AUC) (e.g. Fig. 4a left-hand panel, S26) and standard error of the mean. Significance was determined via ordinary one-way ANOVA using a normal (Gaussian) distribution assumption and multiple comparisons. P values were calculated and graphed using the following designations: P ≤ 0.0001 = **** ; P ≤ 0.001 = *** ; P ≤ 0.01 = ** ; P ≤ 0.05 = * ; P ≥ 0.05 = NS (not significant), by Tukey’s post-hoc test. Cells were imaged live after 4 hours via confocal microscopy utilizing a temperature (37 C) and CO_2_ (5%) regulated chamber. Cells were fixed with 4% PFA after 1 hour and analyzed via flow cytometry.

### Fibroblast/Panc1 co-culture microscopy (Fig. 4d, 4e, S29, S30)

Human dermal fibroblasts were seeded at 5,000 cells/well in a black-walled 96-well glass-bottom tissue culture plate. Panc1 cells were separately seeded at 20,000 cells/well in a 24-well tissue culture plate. After 24 hours, fibroblasts were incubated with Vybrant DiI membrane labeling reagent according to manufacturer protocol (Molecular Probes) for 20 minutes, followed by washing with fresh media to remove unbound labeling reagent. Panc1 cells were incubated with Vybrant DiD membrane labeling reagent for 20 minutes, washed, and trypsinized. The cells were centrifuged and the pellet was redispersed in fibroblast media. Panc1 cells were then seeded at 5,000 cells/well in the same 96-well plate containing fibroblasts to create a co-culture. After 24 hours, media was removed and replaced with 100 µM Rosindol or 10 µM H2D in fibroblast media. Cells were then imaged live via confocal microscopy utilizing a 37° C and 5% CO_2_ regulated chamber after 1 hour to detect H2D fluorescence and over a period of 1 - 4 hours to detect Rosindol fluorescence.

### ROS metabolism inhibitor studies in Panc1 and H6C7 cells (Fig. 4f-4i)

Cells were seeded at 10,000 cells/well in a 96-well black-walled plate. After 24 hours, media was removed and replaced with media containing 100 µM Rosindol, followed by the ROS generation/inhibition control compounds and immediate measurement via microplate reader. 4-octyl itaconate was added 9 hours prior to Rosindol (16 hours after initial plating). Inhibitors were used in the following amounts: rotenone 200 nM, setanaxib 10 µM, LCS-1 10 µM, hydroxylamine 30 µM, brusatol 10 µM, 4-octyl itaconate 75 µM. Each condition was assessed via 3 - 6 replicates per timepoint. After collection, data was imported to Graphpad Prism and plotted as the mean and standard deviation of the replicates at each timepoint (e.g. Fig. S24, S25). Kinetic data was then plotted as area under the curve (AUC) (e.g. Fig. 4g, S26) and standard error of the mean. Significance was determined via ordinary one-way ANOVA using a normal (Gaussian) distribution assumption and multiple comparisons. P values were calculated and graphed using the following designations: P ≤ 0.0001 = **** ; P ≤ 0.001 = *** ; P ≤ 0.01 = ** ; P ≤ 0.05 = * ; P ≥ 0.05 = NS (not significant), by Tukey’s post-hoc test.

### Glucose metabolism study (Fig. 4j)

Panc1, BxPC-3, and H6C7 cells were seeded at 10,000 cells/well in a 96-well black-walled plate. After 24 hours, media was removed and replaced with media containing 100 µM Rosindol, followed by the addition of sufficient β-D-glucose to achieve a final concentration of 200 mM (depending on the initial glucose concentration of each cell media) to a subset of the wells. Fluorescence was then measured via microplate reader over a period of 10 hours. Each condition was assessed via 4 replicates per timepoint. After collection, data was imported to Graphpad Prism and plotted as the mean and standard deviation of the replicates at each timepoint. Kinetic data was then plotted as area under the curve (AUC) and standard error of the mean. Significance was determined via ordinary one-way ANOVA using a normal (Gaussian) distribution assumption and multiple comparisons. P values were calculated and graphed using the following designations: P ≤ 0.0001 = **** ; P ≤ 0.001 = *** ; P ≤ 0.01 = ** ; P ≤ 0.05 = * ; P ≥ 0.05 = NS (not significant), by Tukey’s post-hoc test.

### KRAS^G12D^ and Nrf2 inhibition study (Fig. 4k, 4l, S27)

Panc1 and BxPC-3 cells were seeded at 10,000 cells/well in 96-well black-walled plates. After 15 hours, 100 nM of the KRASG12D inhibitor MRTX1133 was added to a subset of wells, along with 75 µM 4-octyl itaconate in a second subset. After incubation for 9 hours, media was replaced with media containing 100 µM Rosindol, and 10 µM brusatol was added to a third subset of wells. A fourth subset of wells were treated with 100 µM Rosindol and 100 µM H_2_O_2_ to provide a positive control for ROS. Fluorescence was then measured via microplate reader over a period of 10 hours. Each condition was assessed via 4 replicates per timepoint. After collection, data was imported to Graphpad Prism and plotted as the mean and standard deviation of the replicates at each timepoint. Kinetic data was then plotted as area under the curve (AUC) and standard error of the mean. The AUC for cells treated with Rosindol in conjunction with MRTX1133, 4-octyl itaconate, and brusatol were divided by the AUC of cells treated with Rosindol alone to determine the % change in basal ROS signal in the presence of each small molecule (Fig. 4k, 4l). Significance was determined via ordinary one-way ANOVA using a normal (Gaussian) distribution assumption and multiple comparisons. P values were calculated and graphed using the following designations: P ≤ 0.0001 = **** ; P ≤ 0.001 = *** ; P ≤ 0.01 = ** ; P ≤ 0.05 = * ; P ≥ 0.05 = NS (not significant), by Tukey’s post-hoc test.

## Availability of data and materials

The raw data required to reproduce these findings are available from the authors upon request.

## Author Contributions

Conceptualization: AWM

Methodology: AWM, MNN

Investigation: AWM, MNN, CDS, JJM, NA, AL

Visualization: AWM

Supervision: MWG

Writing—original draft: AWM

Writing—review & editing: All authors Funding: MWG

## Acknowledgements

We acknowledge and thank the Grace Yang and Heather Murray for help.

## Supplementary Information

**Figure S1.**
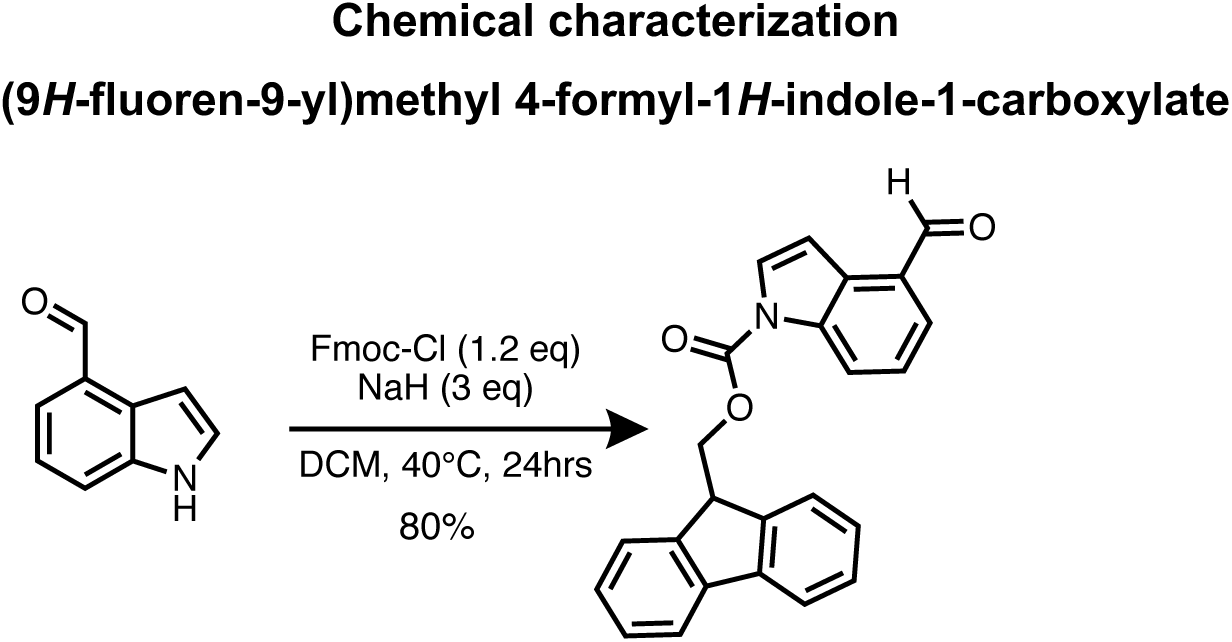
Synthetic scheme of (9*H*-fluoren-9-yl)methyl 4-formyl-1*H*-indole-1-carboxylate.

**Figure S2.**
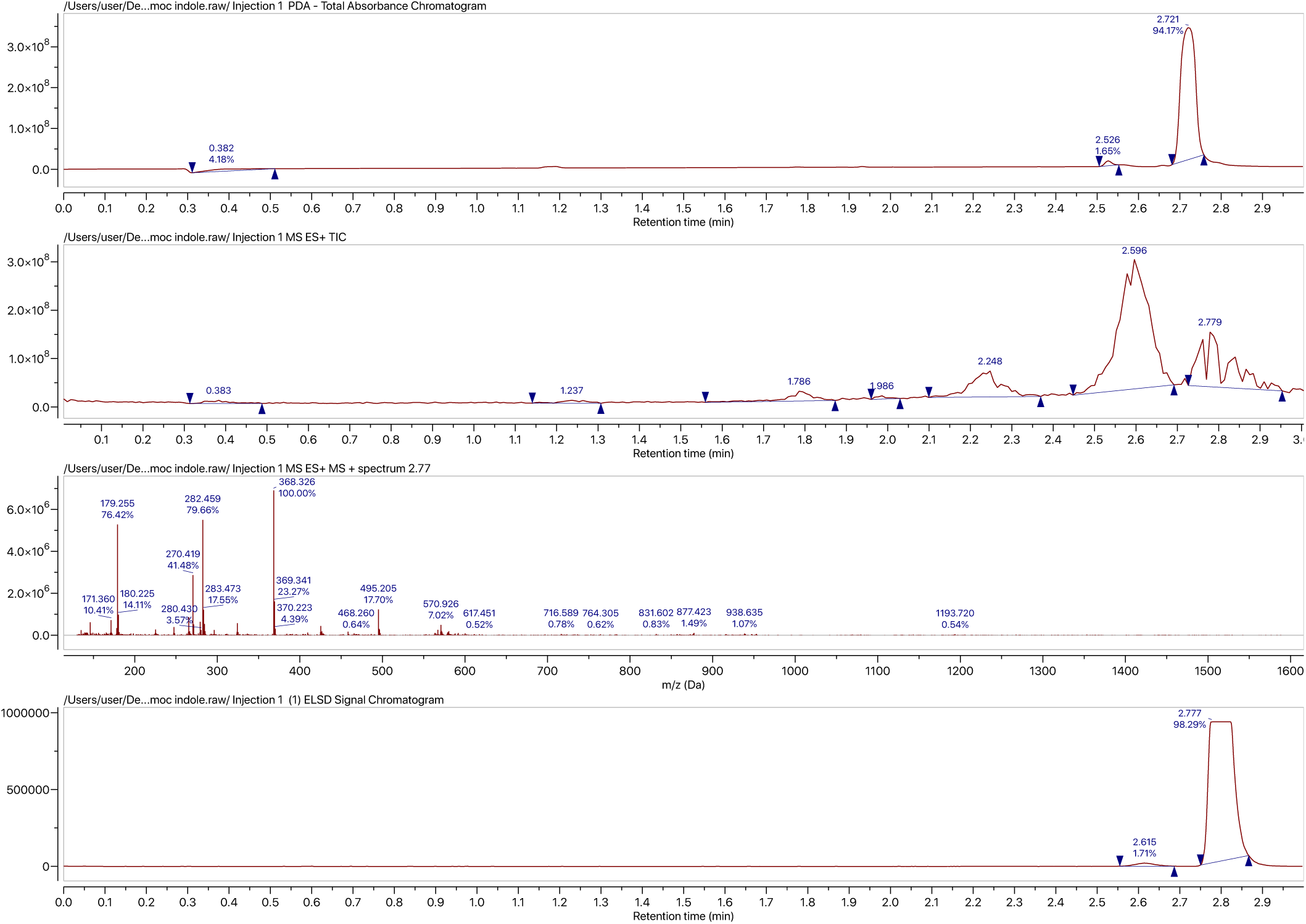
UPLC-MS spectrum of (9*H*-fluoren-9-yl)methyl 4-formyl-1*H*-indole-1-carboxylate.

**Figure S3.**
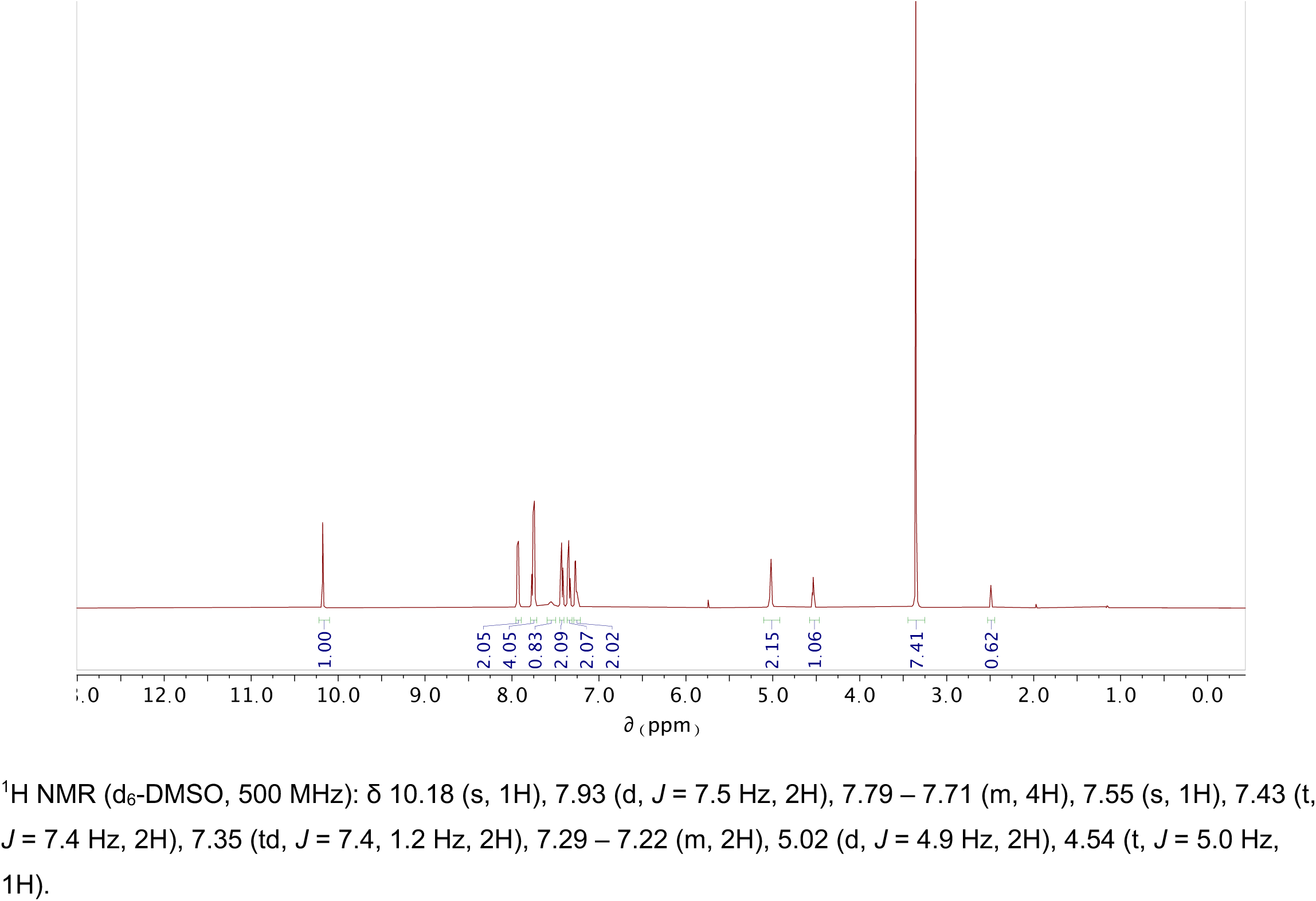
^1^H-NMR spectrum of (9*H*-fluoren-9-yl)methyl 4-formyl-1*H*-indole-1-carboxylate.

**Figure S4.**
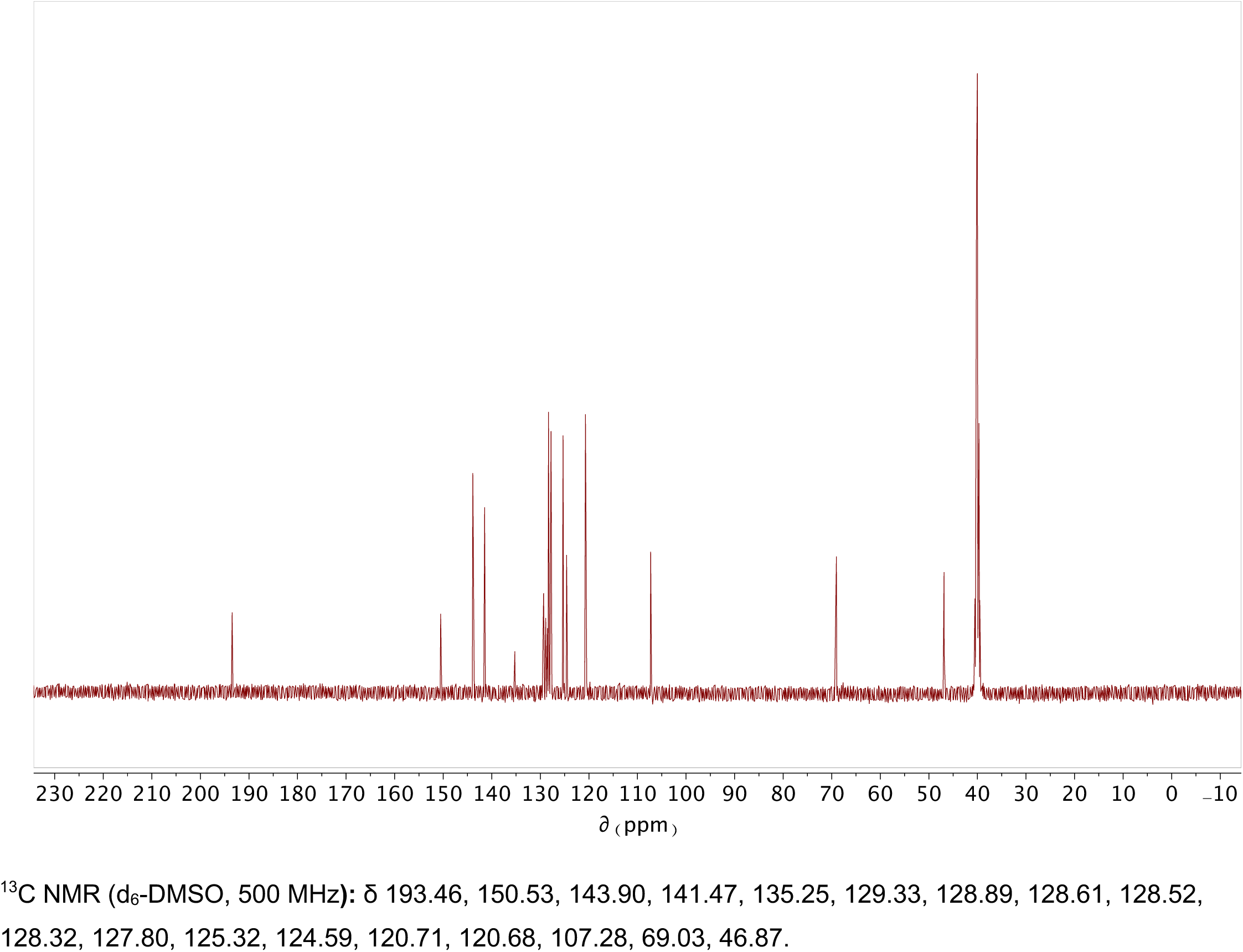
^13^C-NMR spectrum of (9*H*-fluoren-9-yl)methyl 4-formyl-1*H*-indole-1-carboxylate.

**Figure S5.**
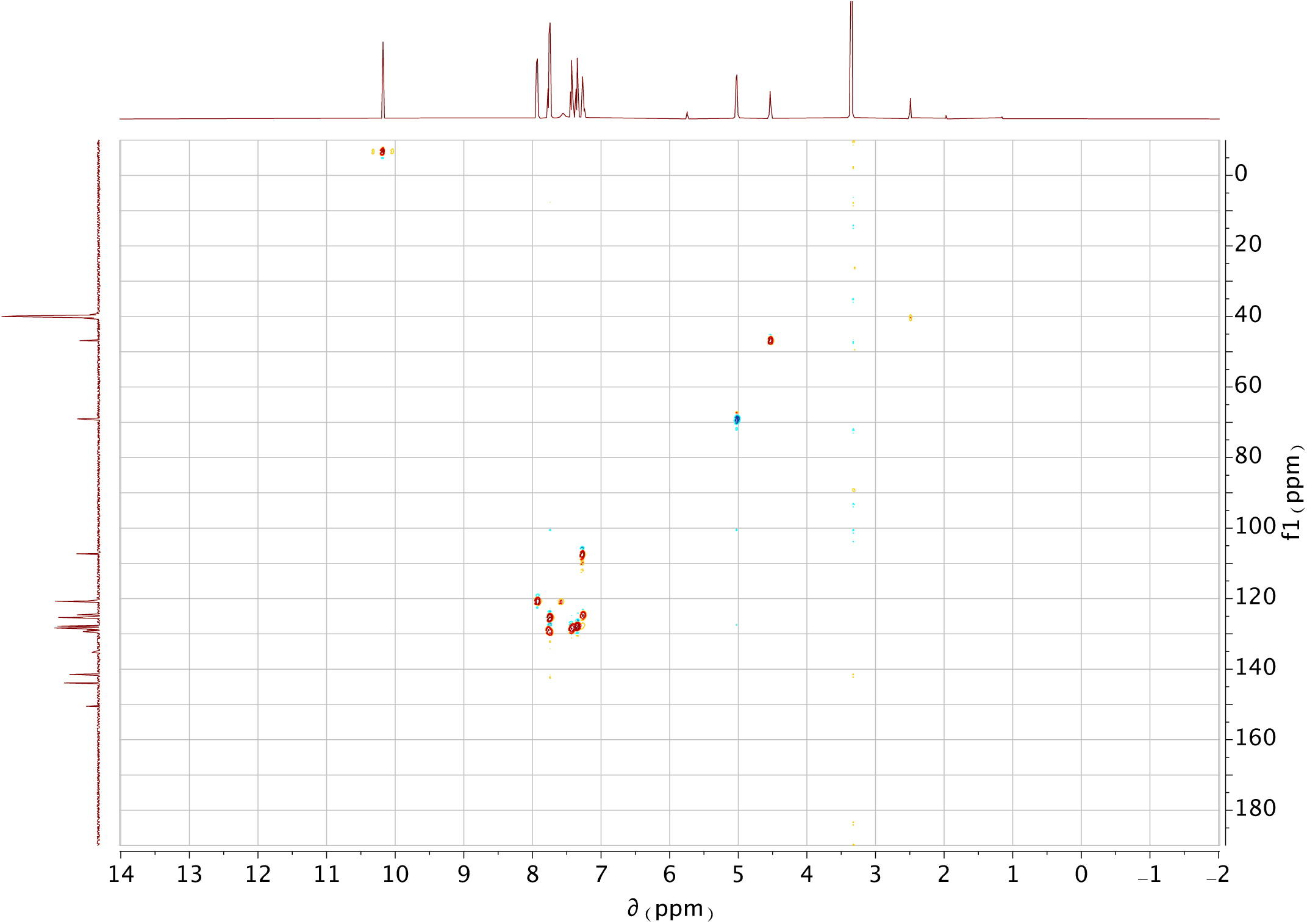
H/C HSQC-NMR spectrum of (9*H*-fluoren-9-yl)methyl 4-formyl-1*H*-indole-1-carboxylate.

**Figure S6.**
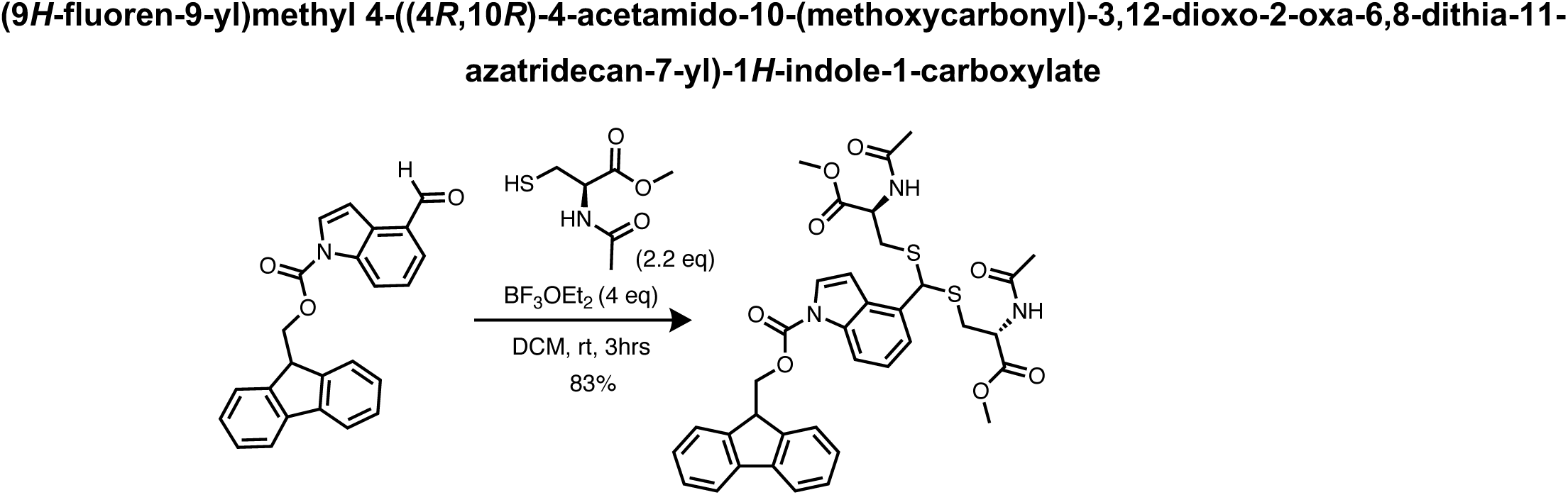
Synthetic scheme of (9*H*-fluoren-9-yl)methyl 4-((4*R*,10*R*)-4-acetamido-10-(methoxycarbonyl)-3,12-dioxo-2-oxa-6,8-dithia-11-azatridecan-7-yl)-1*H*-indole-1-carboxylate.

**Figure S7.**
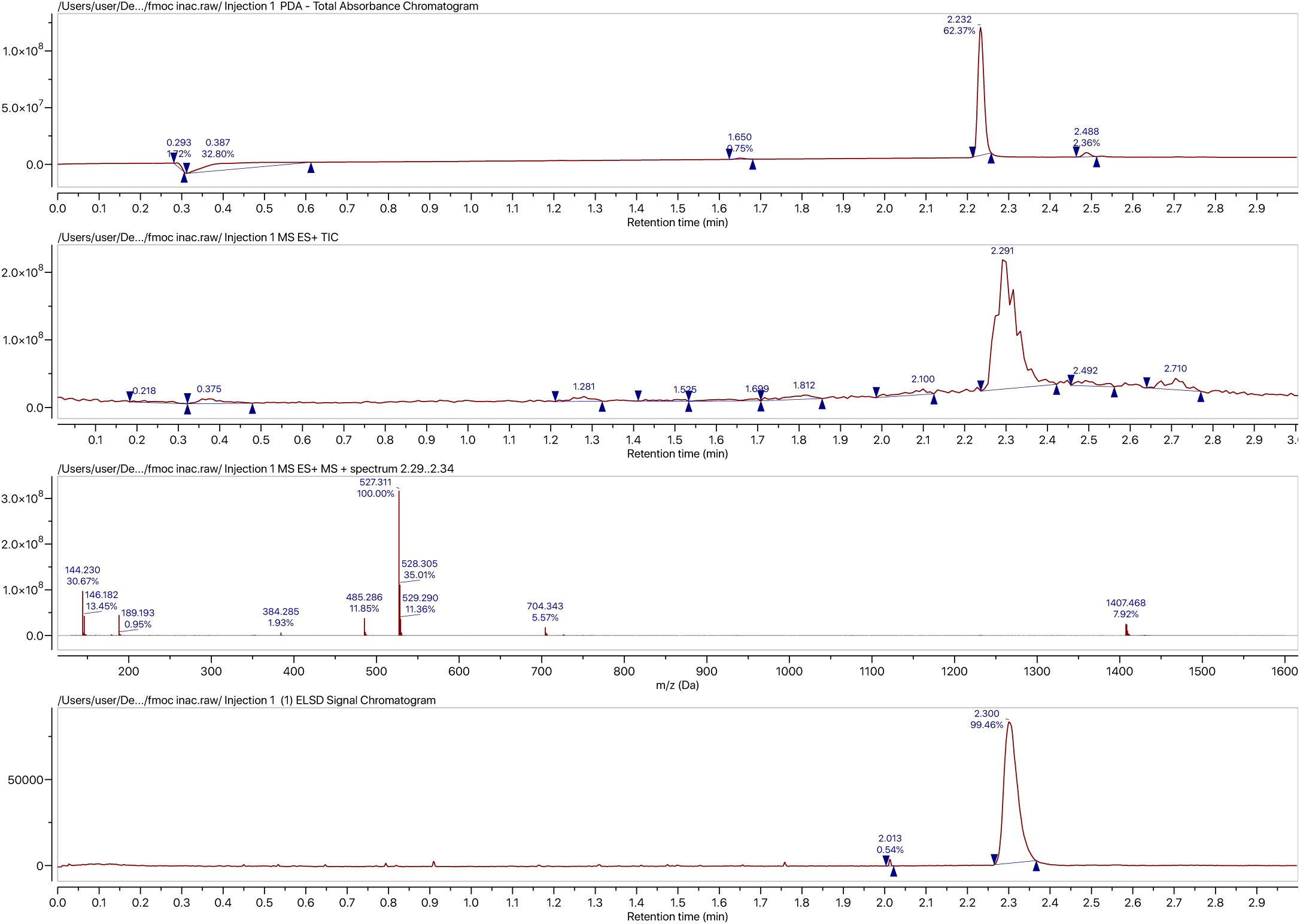
UPLC-MS spectrum of (9*H*-fluoren-9-yl)methyl 4-((4*R*,10*R*)-4-acetamido-10-(methoxycarbonyl)-3,12-dioxo-2-oxa-6,8-dithia-11-azatridecan-7-yl)-1*H*-indole-1-carboxylate.

**Figure S8.**
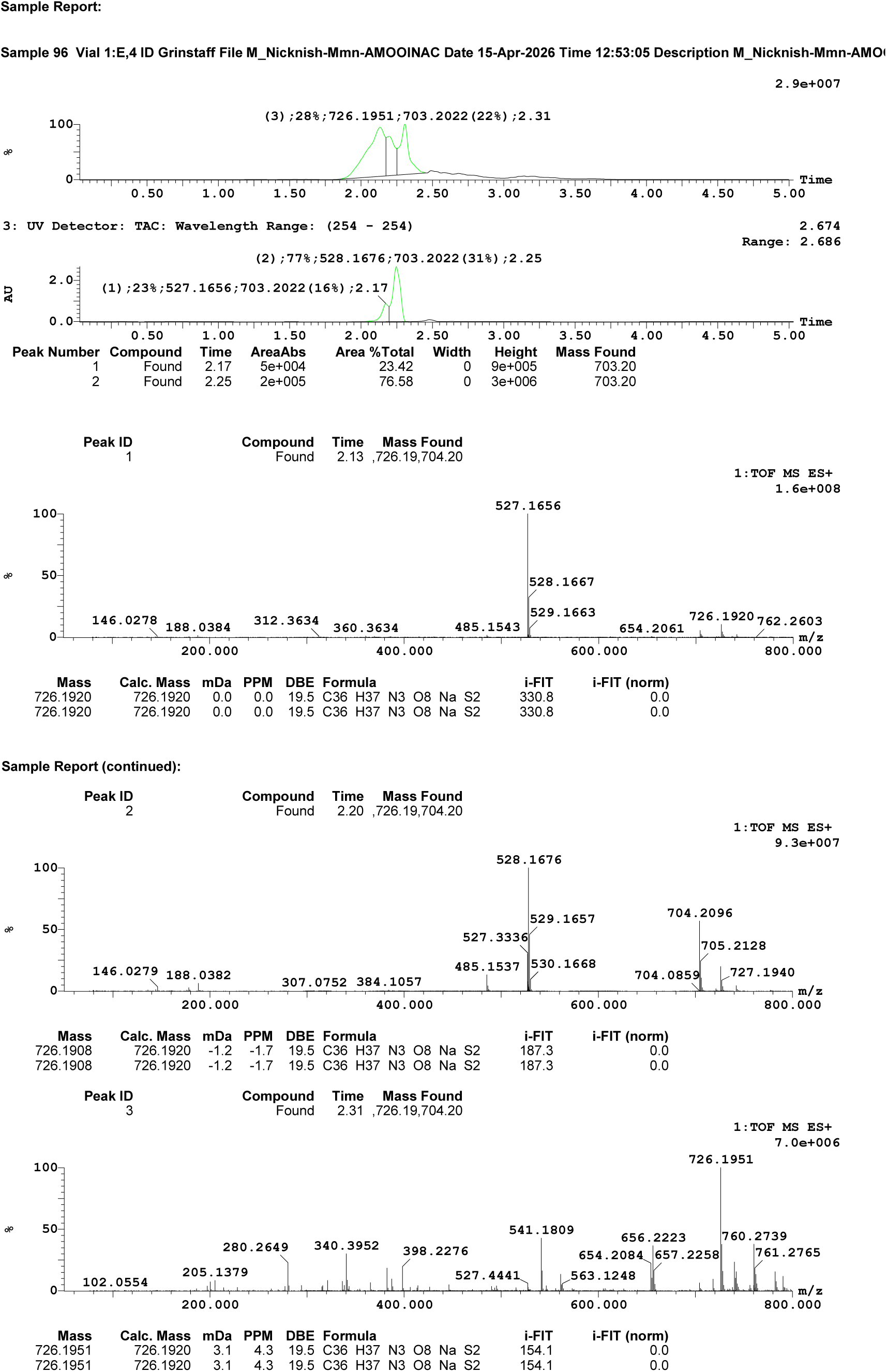
High resolution-MS spectrum of (9*H*-fluoren-9-yl)methyl 4-((4*R*,10*R*)-4-acetamido-10-(methoxycarbonyl)-3,12-dioxo-2-oxa-6,8-dithia-11-azatridecan-7-yl)-1*H*-indole-1-carboxylate.

**Figure S9.**
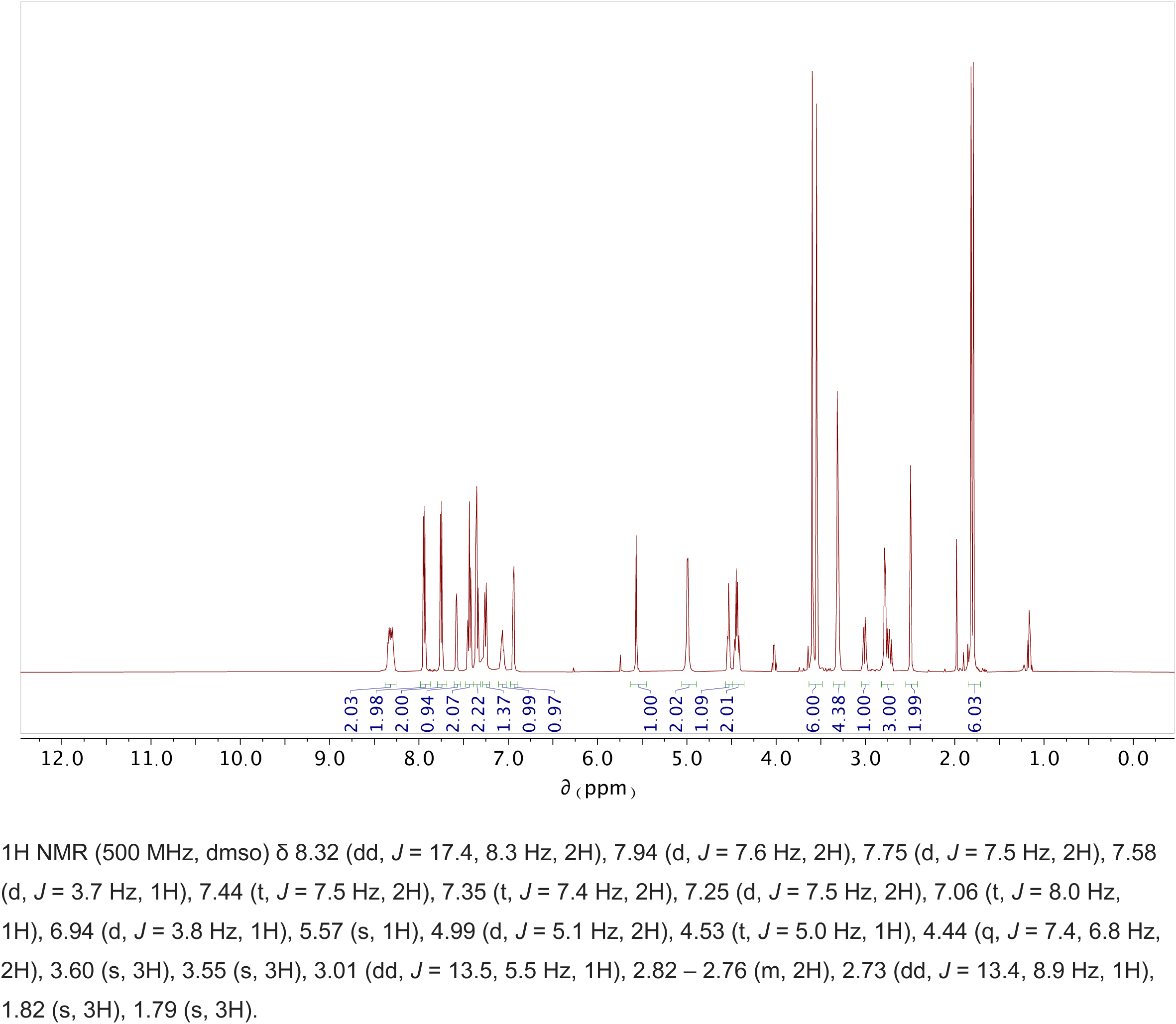
**^1^**H-NMR spectrum of (9*H*-fluoren-9-yl)methyl 4-((4*R*,10*R*)-4-acetamido-10-(methoxycarbonyl)-3,12-dioxo-2-oxa-6,8-dithia-11-azatridecan-7-yl)-1*H*-indole-1-carboxylate.

**Figure S10.**
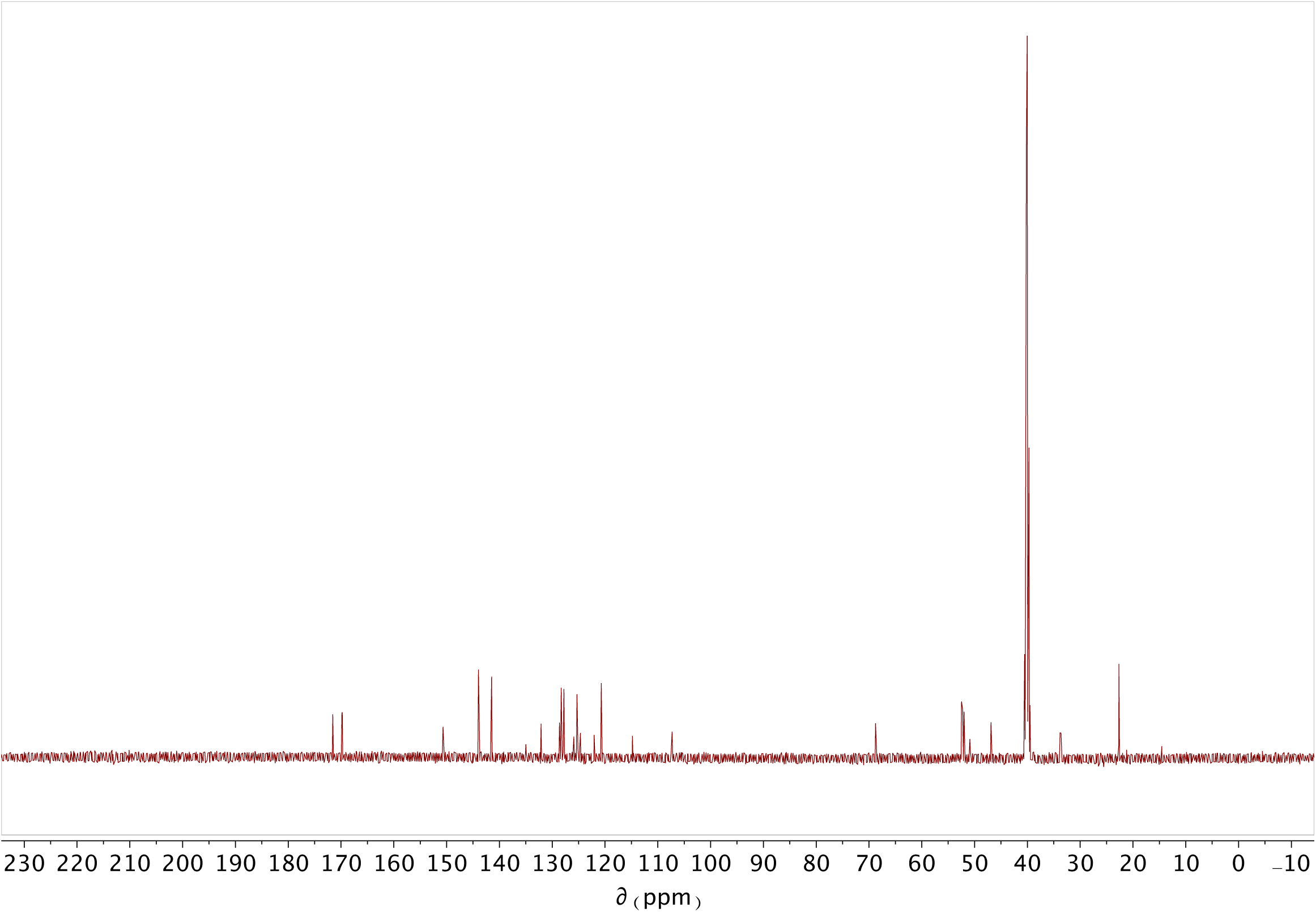
**^13^**C-NMR spectrum of (9*H*-fluoren-9-yl)methyl 4-((4*R*,10*R*)-4-acetamido-10-(methoxycarbonyl)-3,12-dioxo-2-oxa-6,8-dithia-11-azatridecan-7-yl)-1*H*-indole-1-carboxylate.

**Figure S11.**
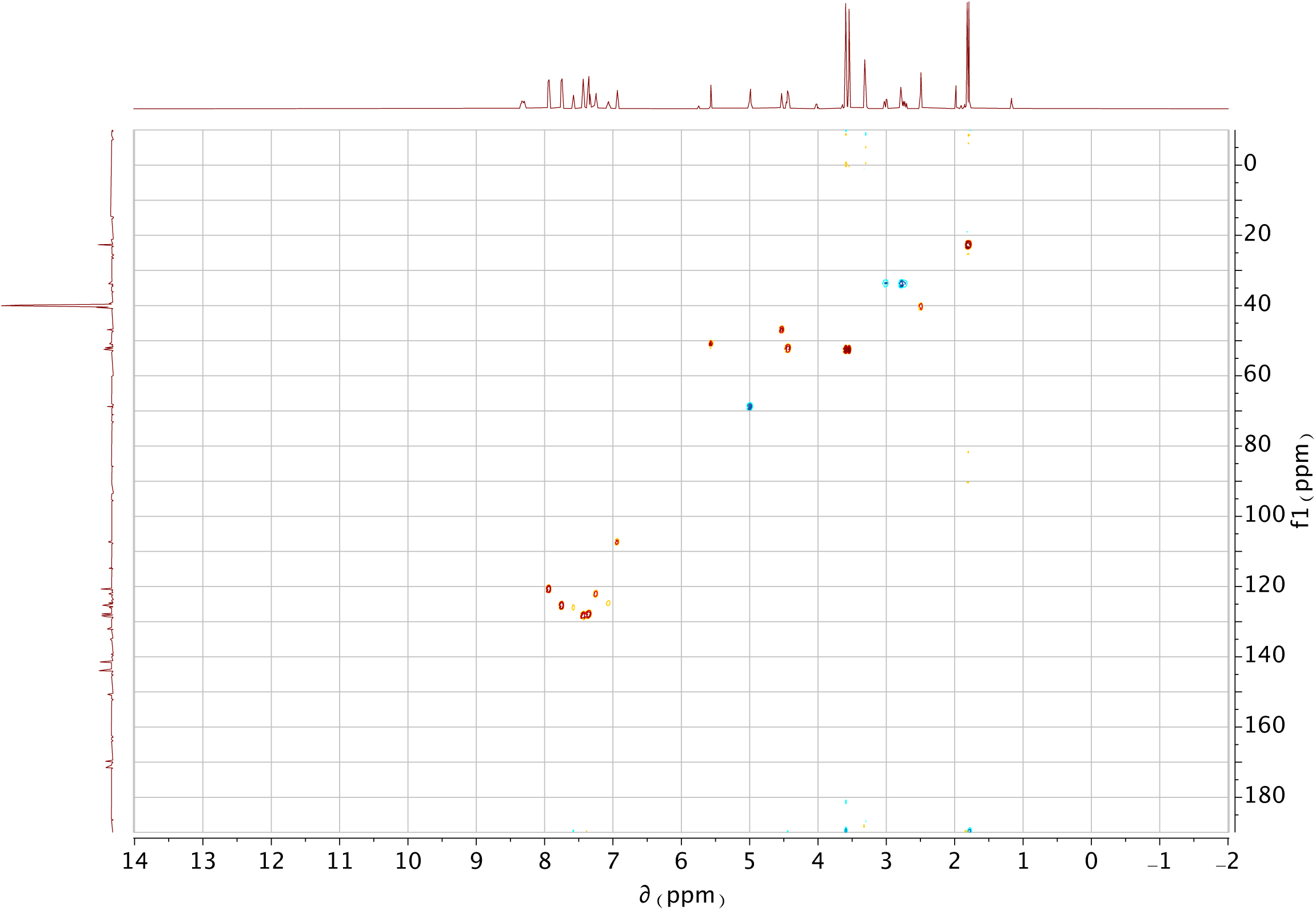
H/C HSQC-NMR spectrum of (9*H*-fluoren-9-yl)methyl 4-((4*R*,10*R*)-4-acetamido-10-(methoxycarbonyl)-3,12-dioxo-2-oxa-6,8-dithia-11-azatridecan-7-yl)-1*H*-indole-1-carboxylate.

**Figure S12.**
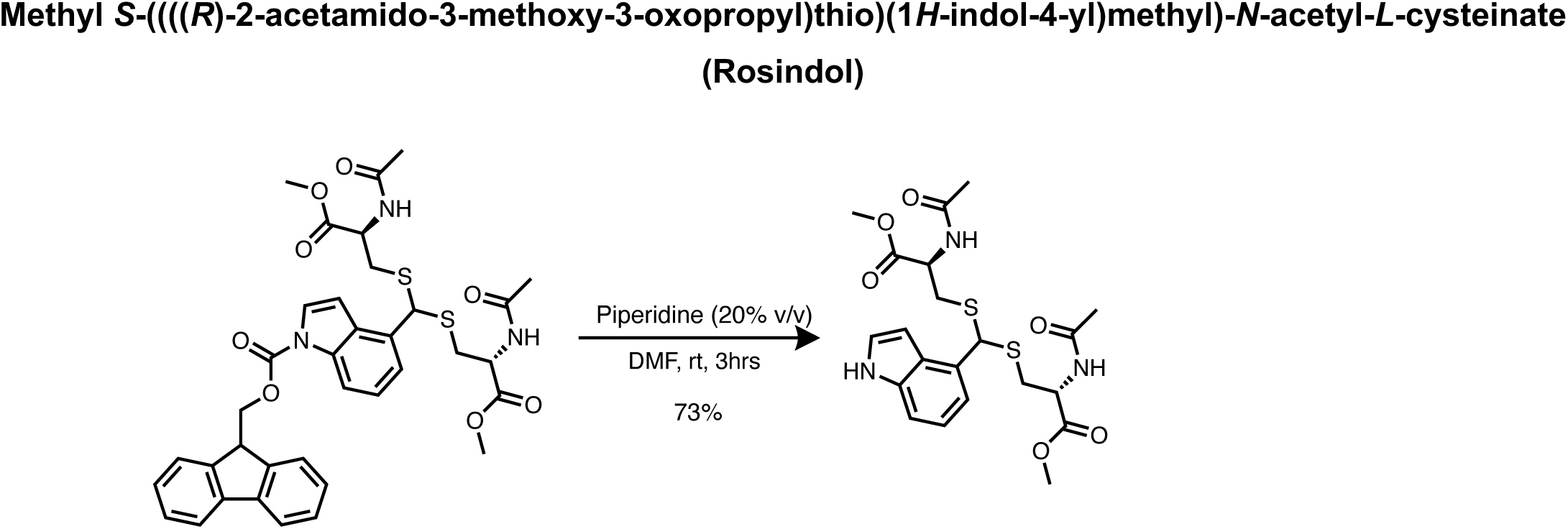
Synthetic scheme of Methyl *S*-((((*R*)-2-acetamido-3-methoxy-3-oxopropyl)thio)(1*H*-indol-4-yl)methyl)-*N*-acetyl-*L*-cysteinate (Rosindol)

**Figure S13.**
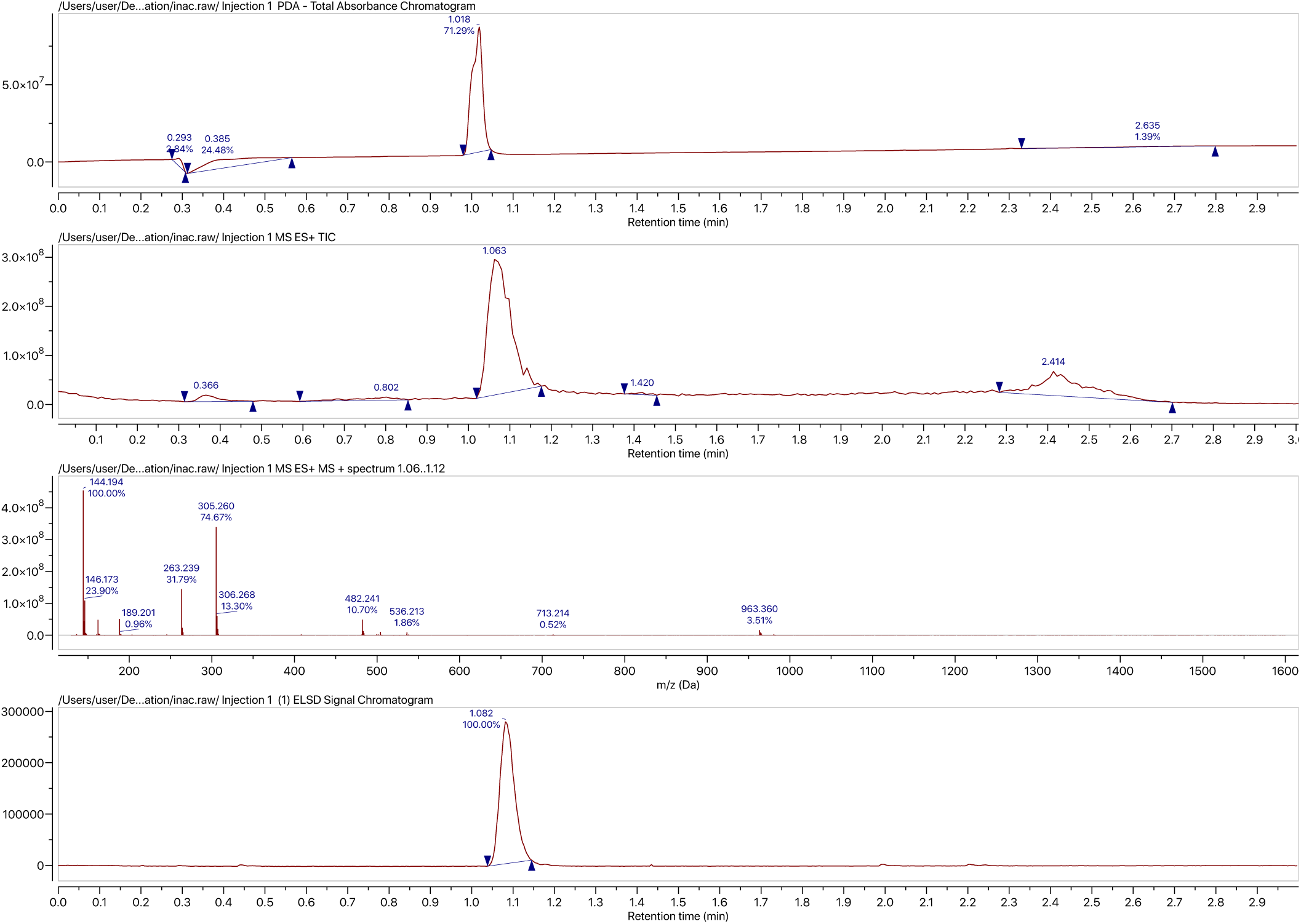
UPLC-MS spectrum of Rosindol.

**Figure S14.**
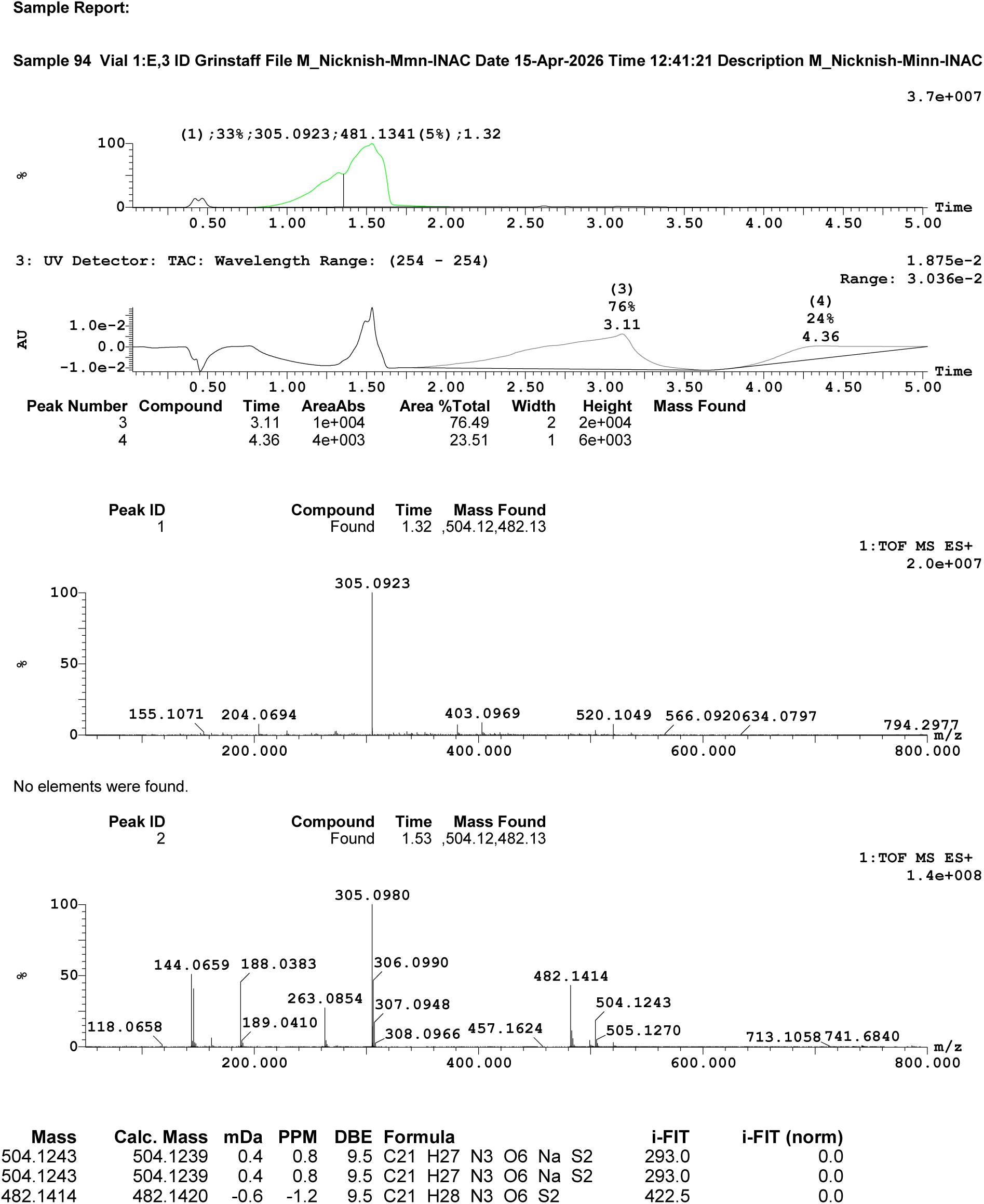
High resolution MS spectrum of Rosindol.

**Figure S15.**
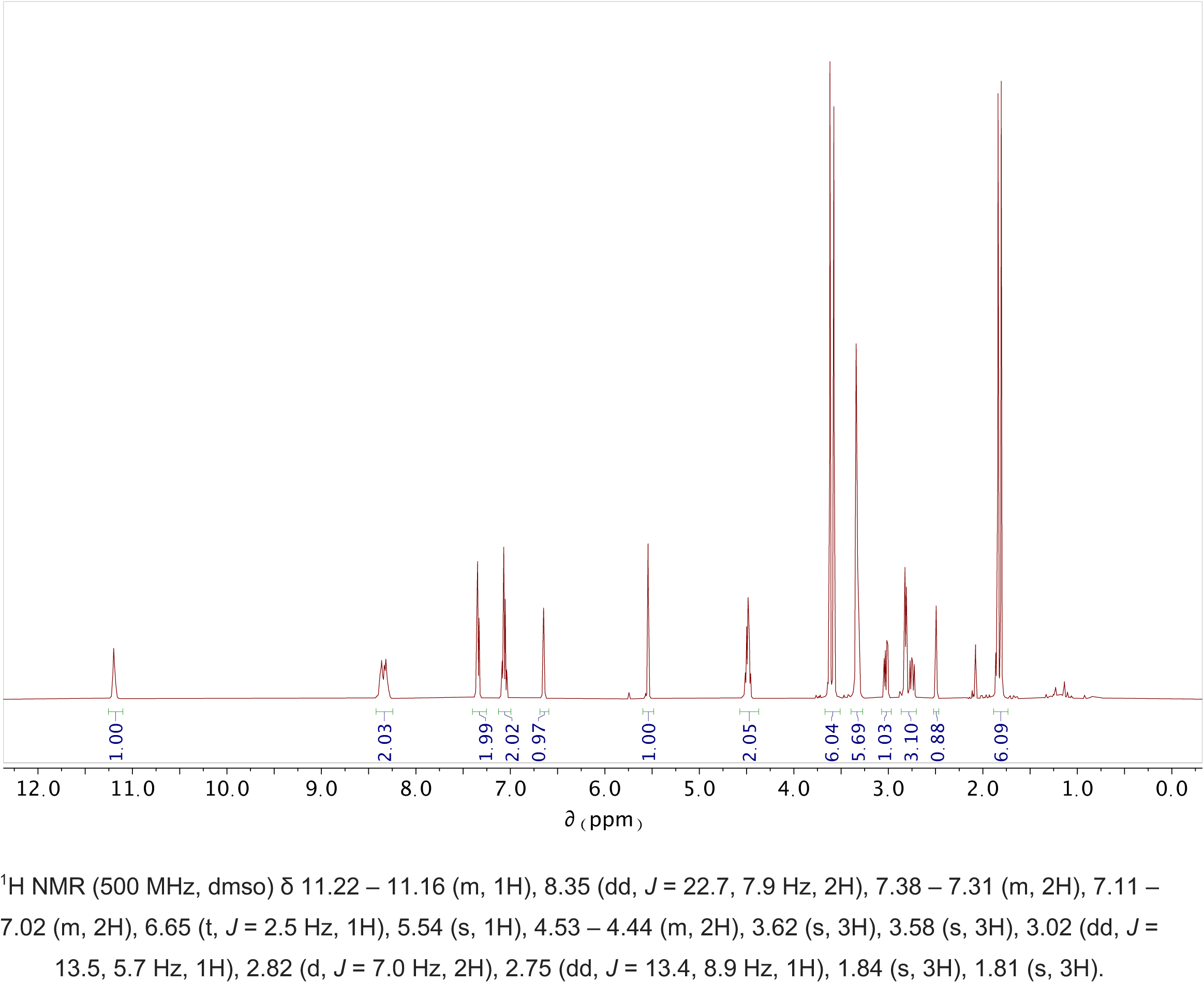
^1^H-NMR spectrum of Rosindol.

**Figure S16.**
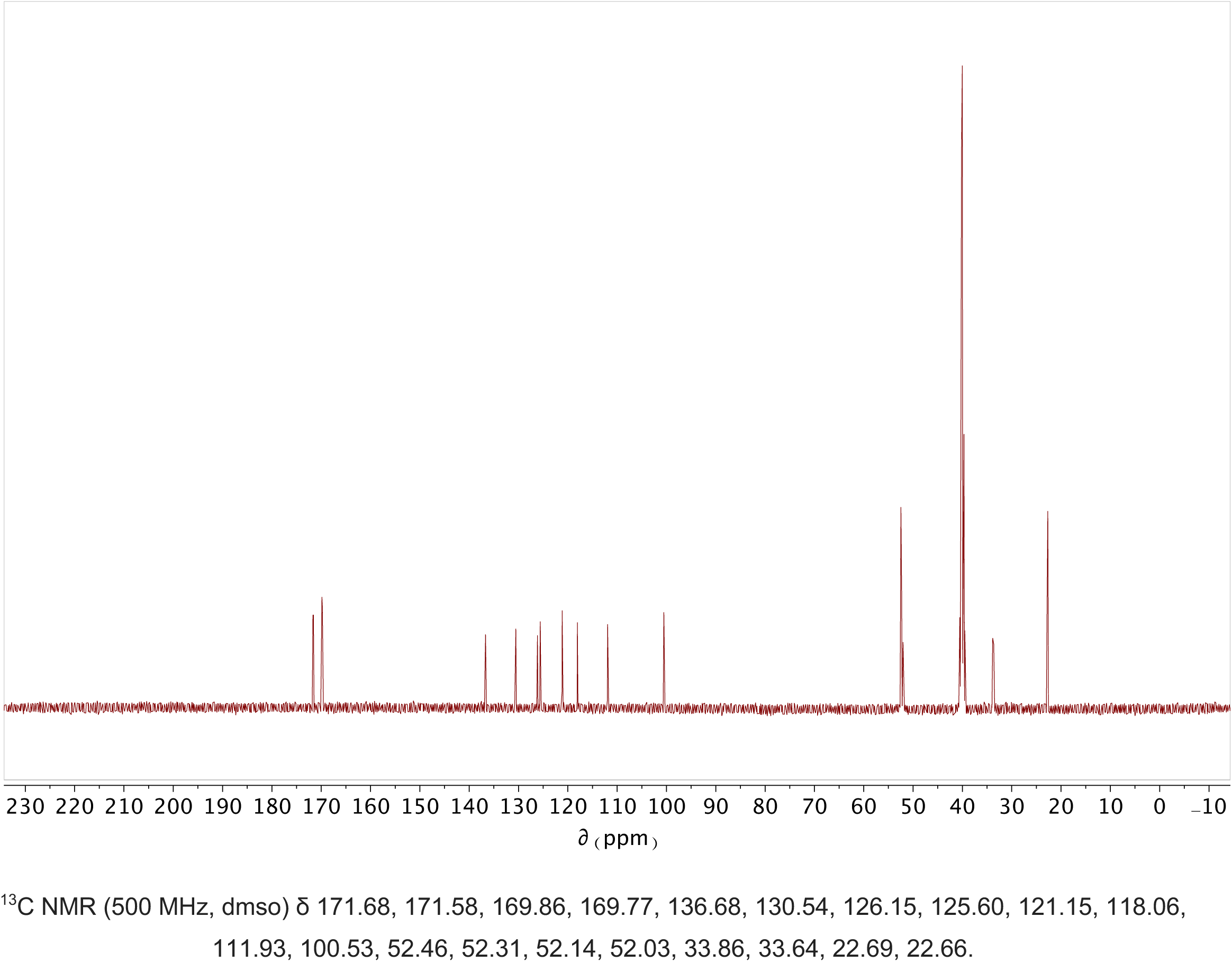
^13^C-NMR spectrum of Rosindol.

**Figure S17.**
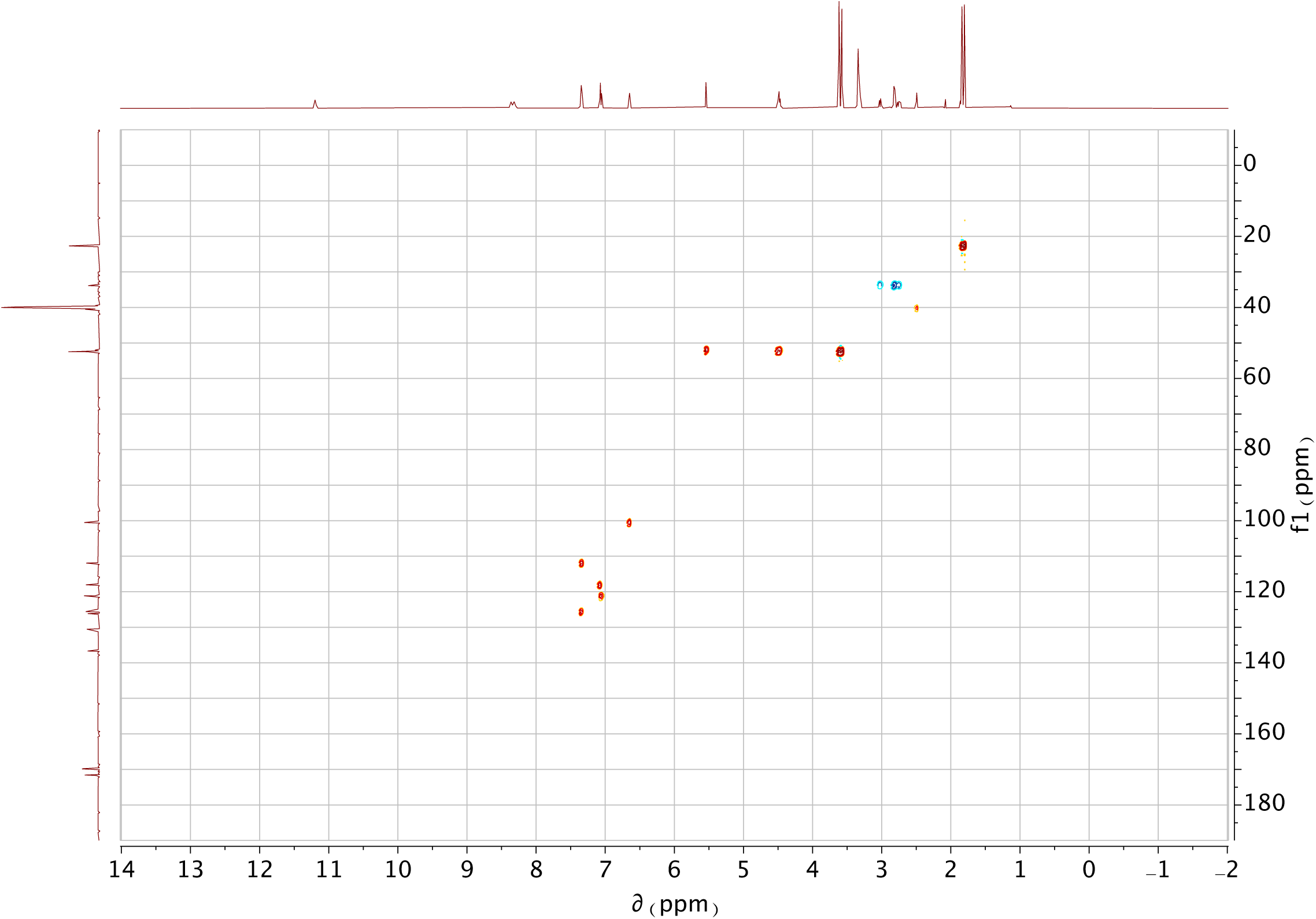
H/C HSQC-NMR spectrum of Rosindol.

**Figure S18.**
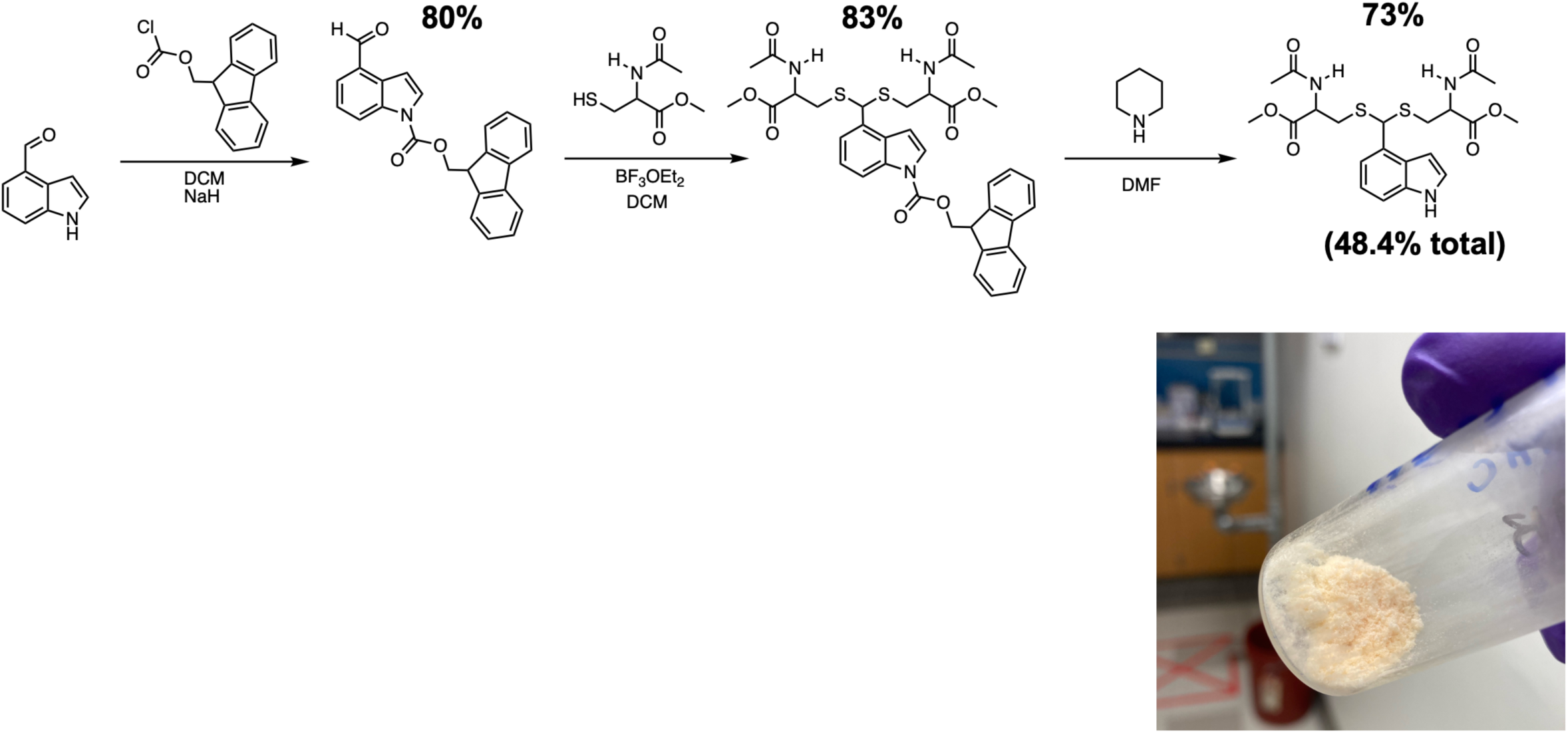
Synthetic route, yields, and gross visual appearance of Rosindol.

**Figure S19.**
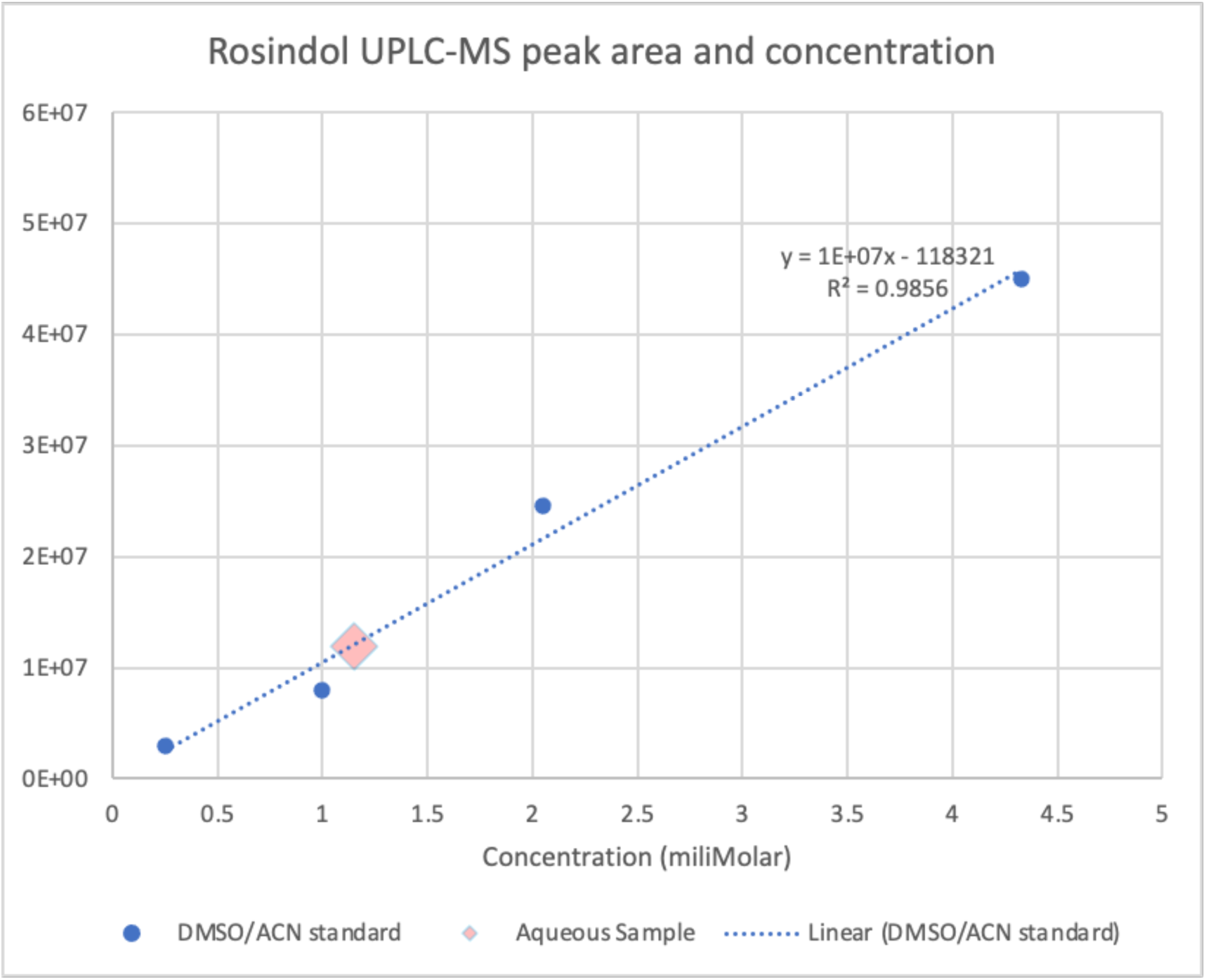
Aqueous solubility of Rosindol via UPLC-MS peak area.

**Figure S20.**
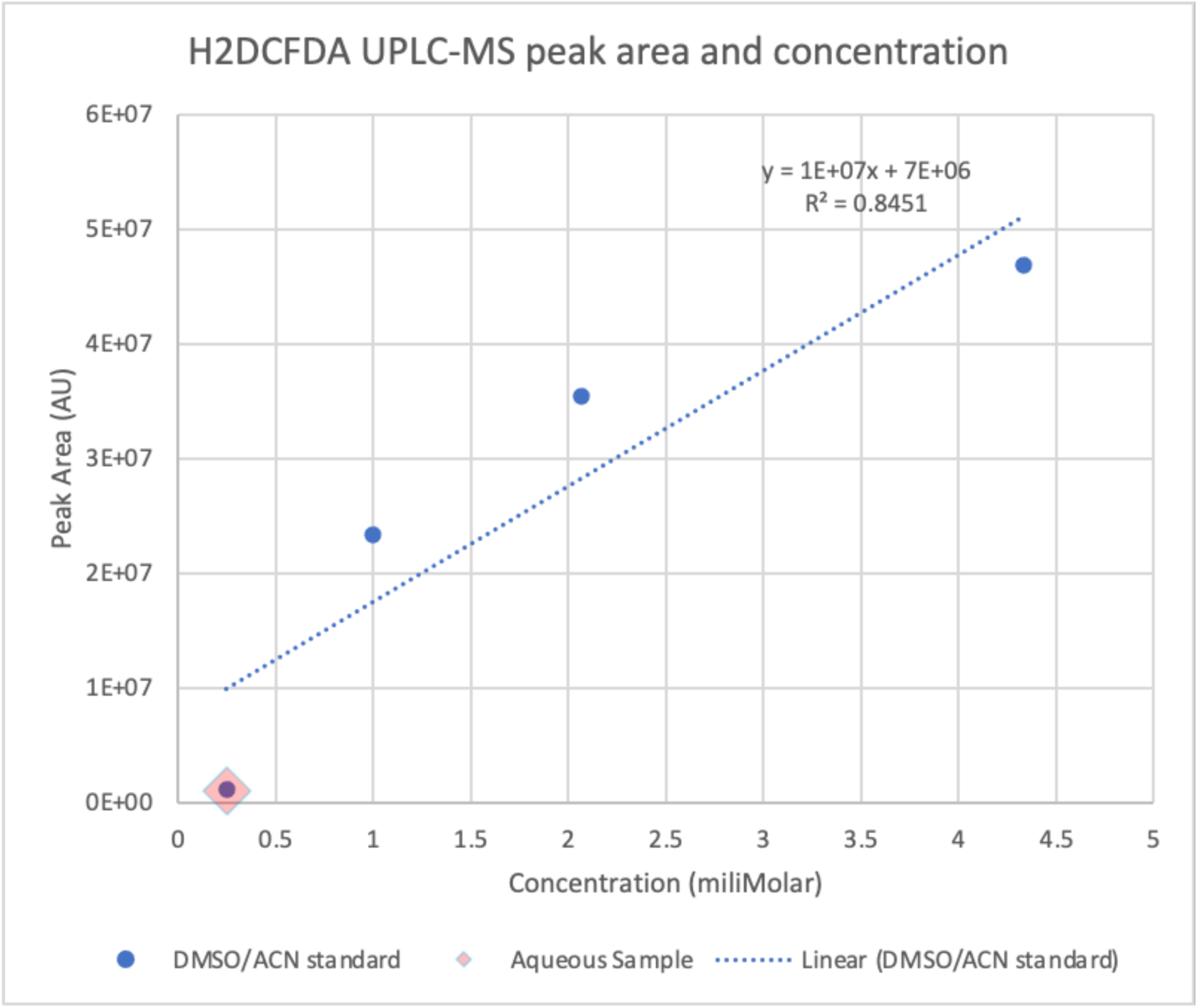
Aqueous solubility of H2D via UPLC-MS peak area.

**Figure S21.**
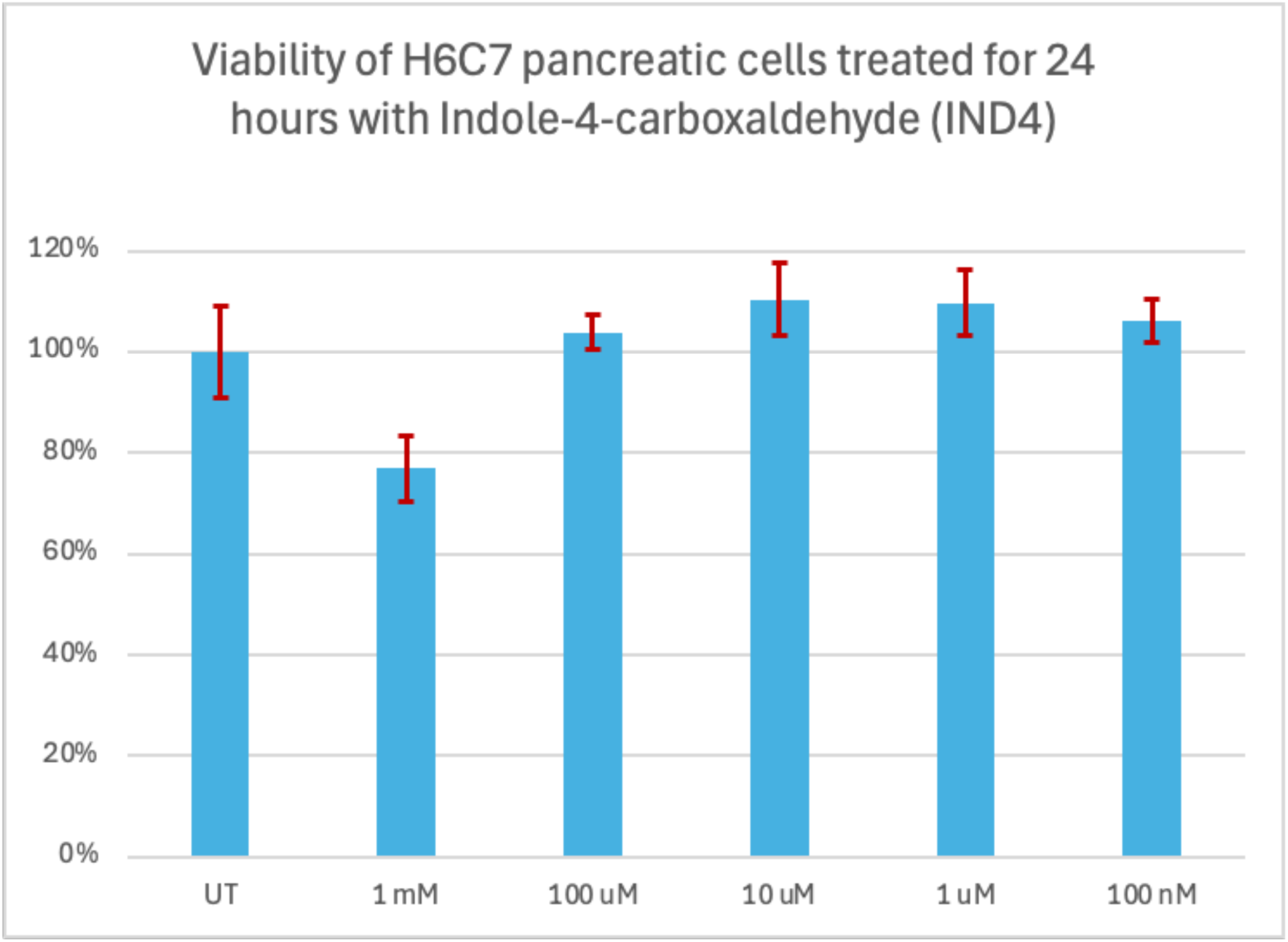
Cytotoxicity of indole-4-carboxaldehyde (IND4) in H6C7 cells via MTS assay.

**Figure S22.**
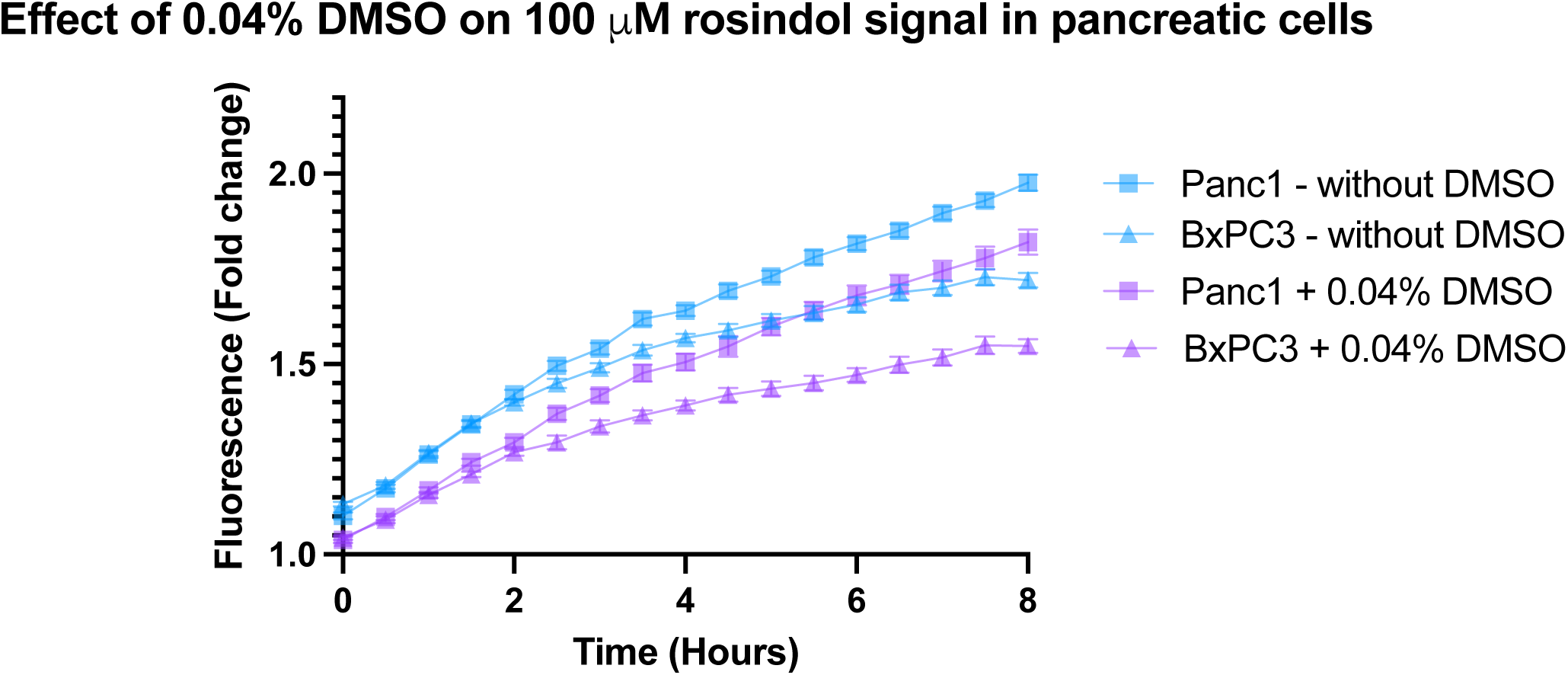
ROS measurements in Panc1 and BxPC-3 cells with and without 0.04% DMSO as added cosolvent. Aqueous Rosindol without DMSO was prepared via adding 0.63 mg of Rosindol to 13 mL of cell media in a conical tube and sonicating for 5 minutes.

**Figure S23.**
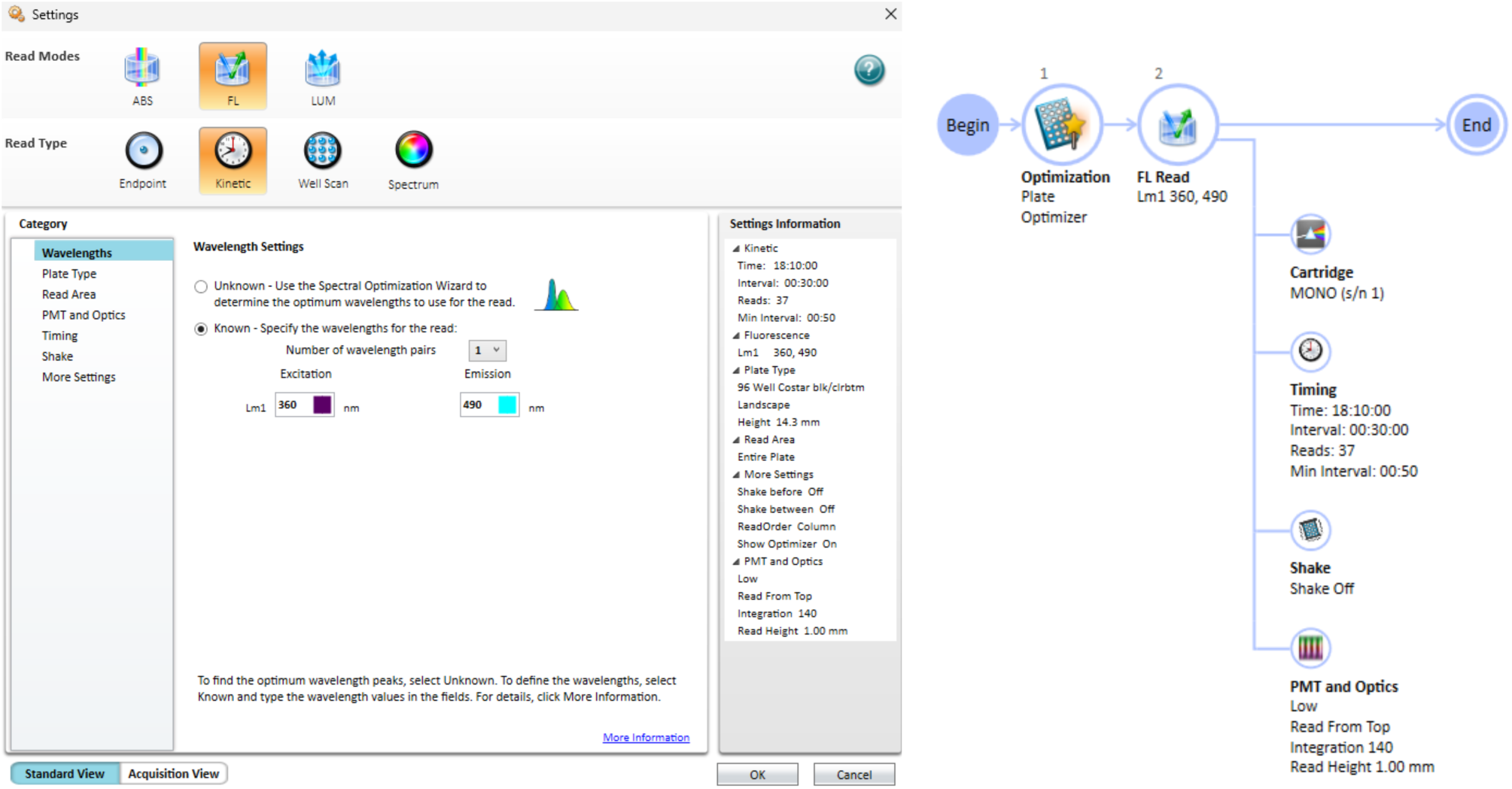
Sample instrument settings in Softmax Pro 7 for live cell kinetic fluorescence measurements via microplate reader.

**Figure S24.**
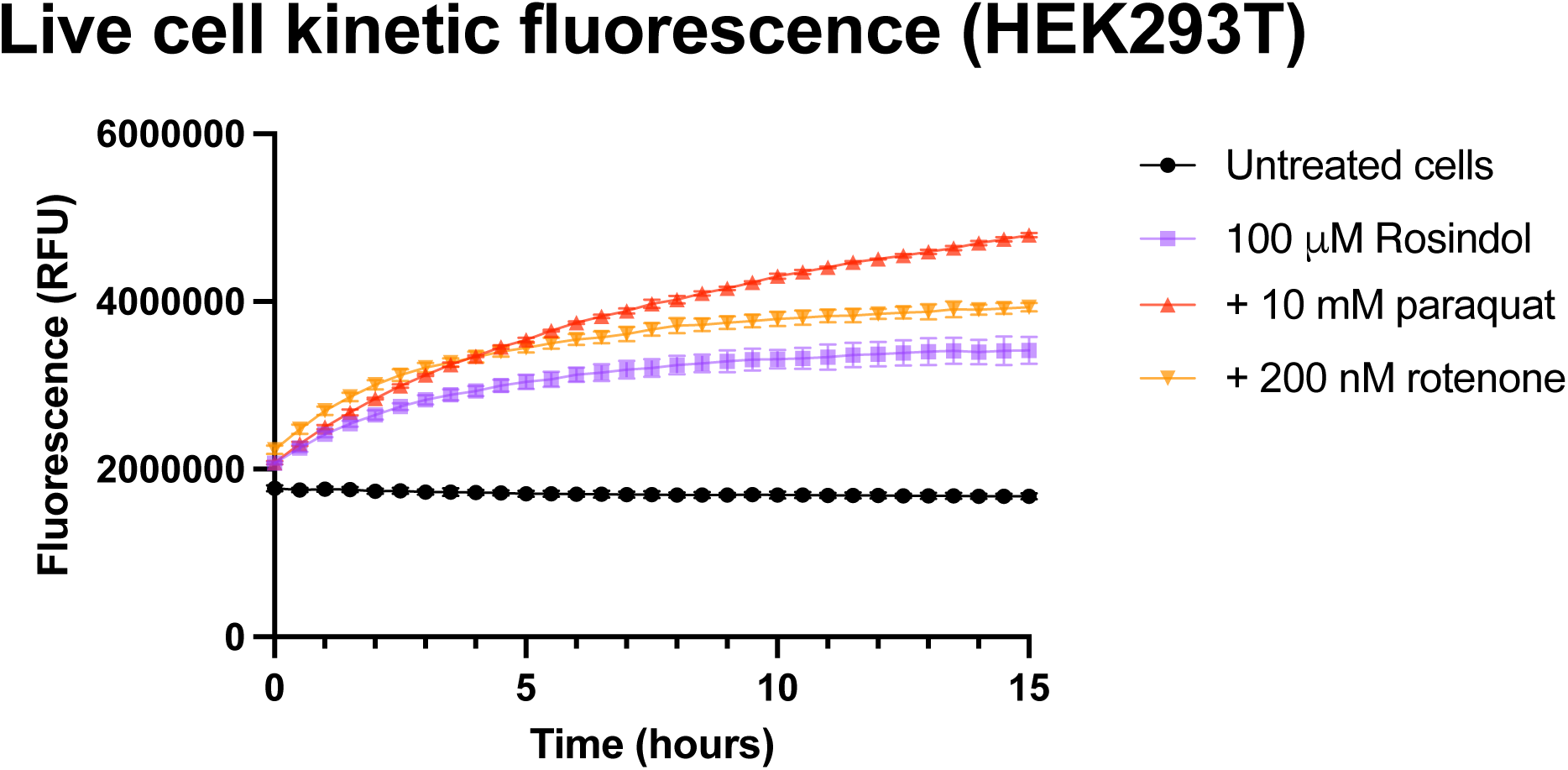
Sample measurements collected from live cell kinetic fluorescence experiment via microplate reader. Each data point reflects the mean of 4 replicates, and error bars represent the standard deviation.

**Figure S25.**
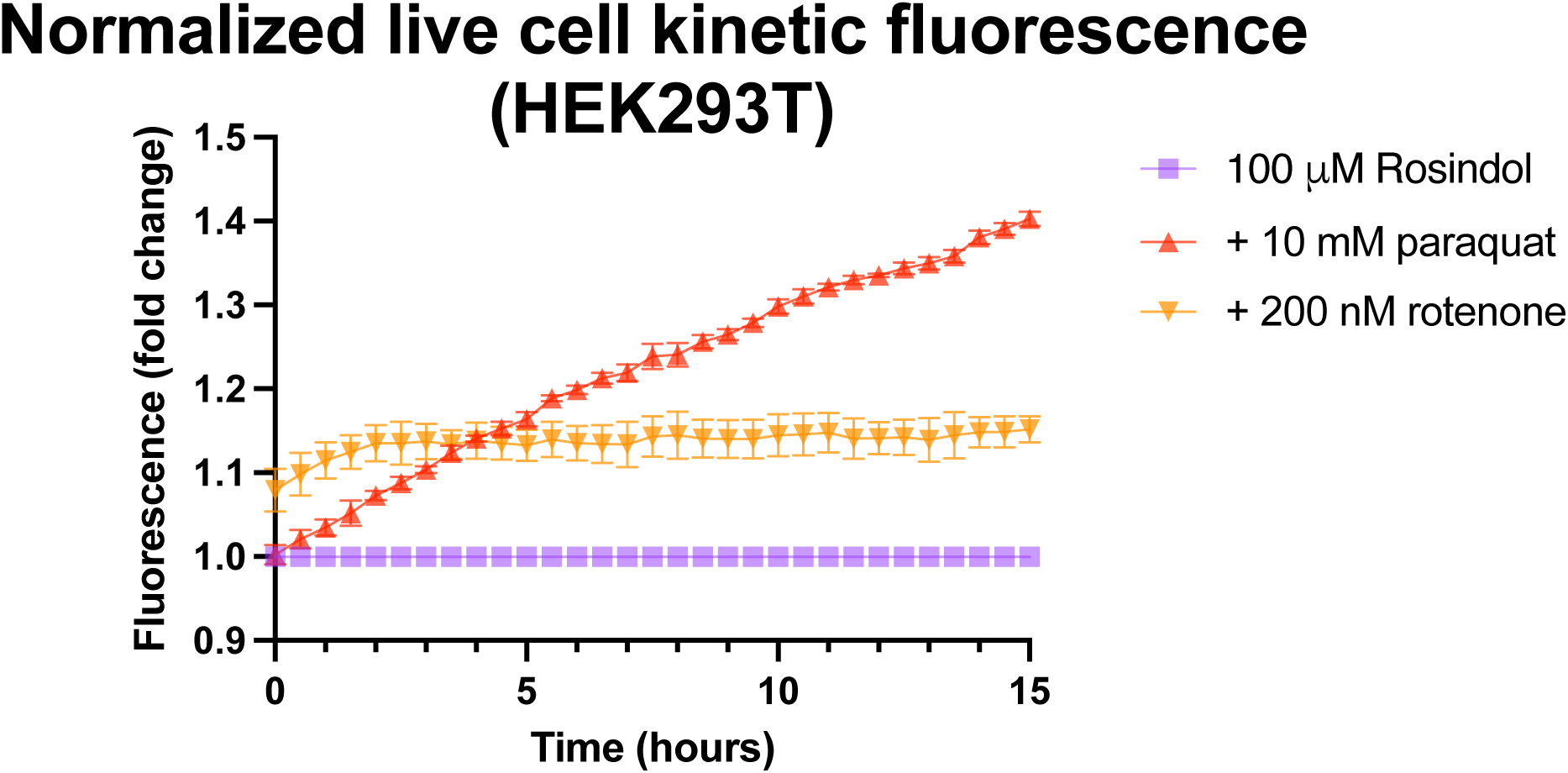
Normalized sample measurements after dividing the magnitude of each data point in Fig. S24 by the magnitude of the Rosindol signal (untreated cells omitted).

**Figure S26.**
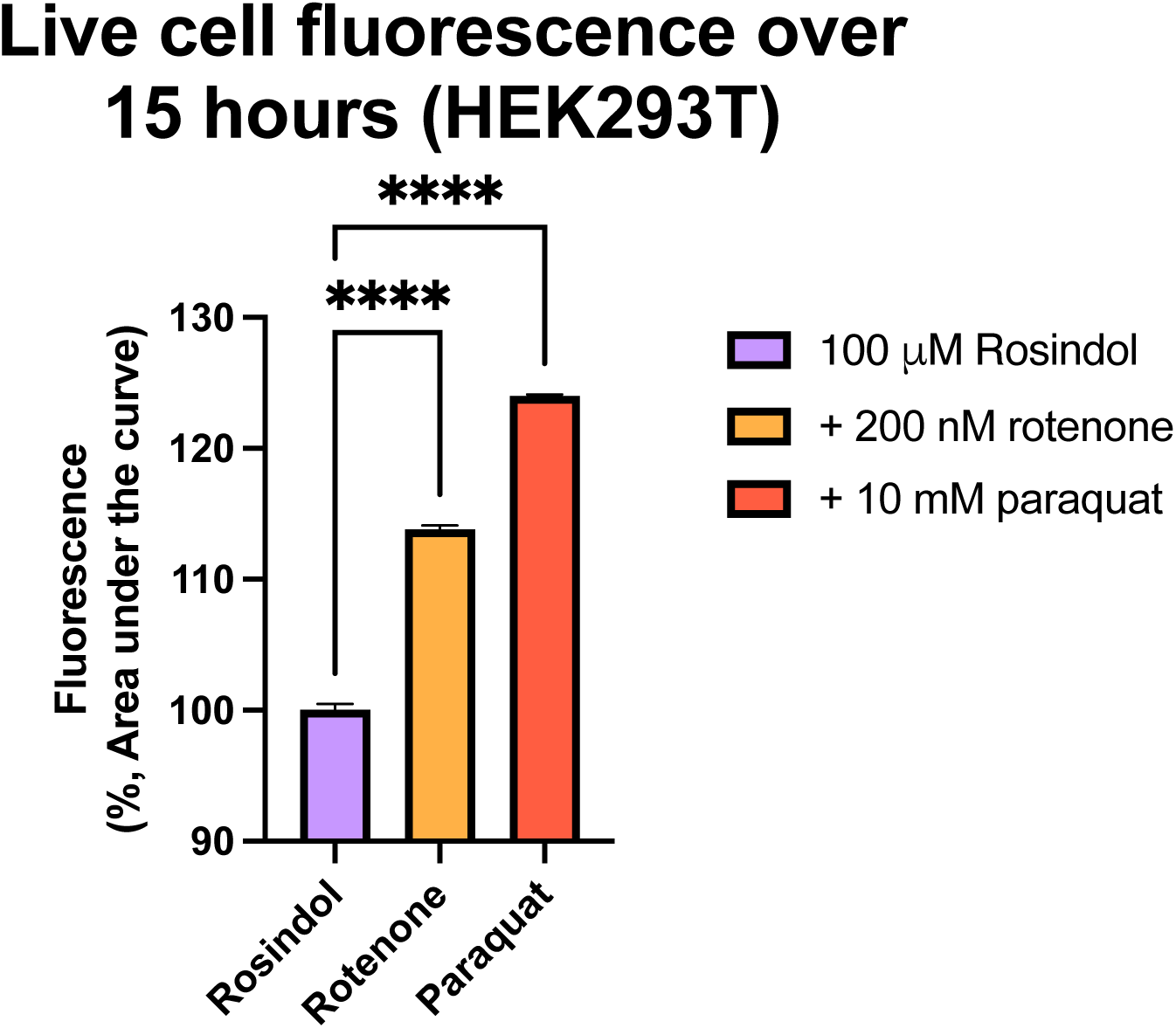
Sample measurements after area performing under the curve (AUC) analysis on kinetic data in Fig. S24, with significance determined via one-way ANOVA.

**Figure S27.**
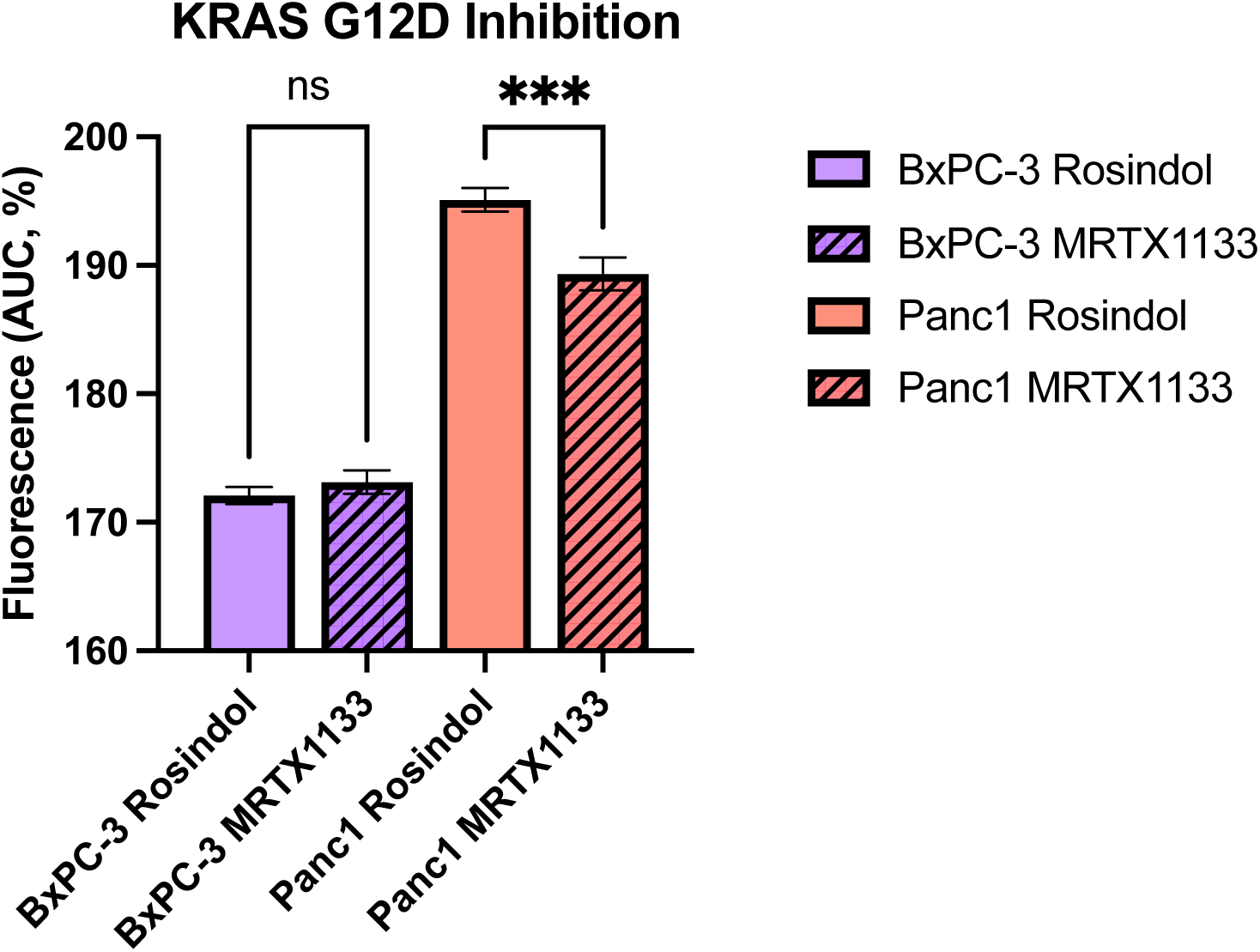
Fluorescence of Panc1 and BxPC-3 cells with and without MRTX1133.

**Figure S28.**
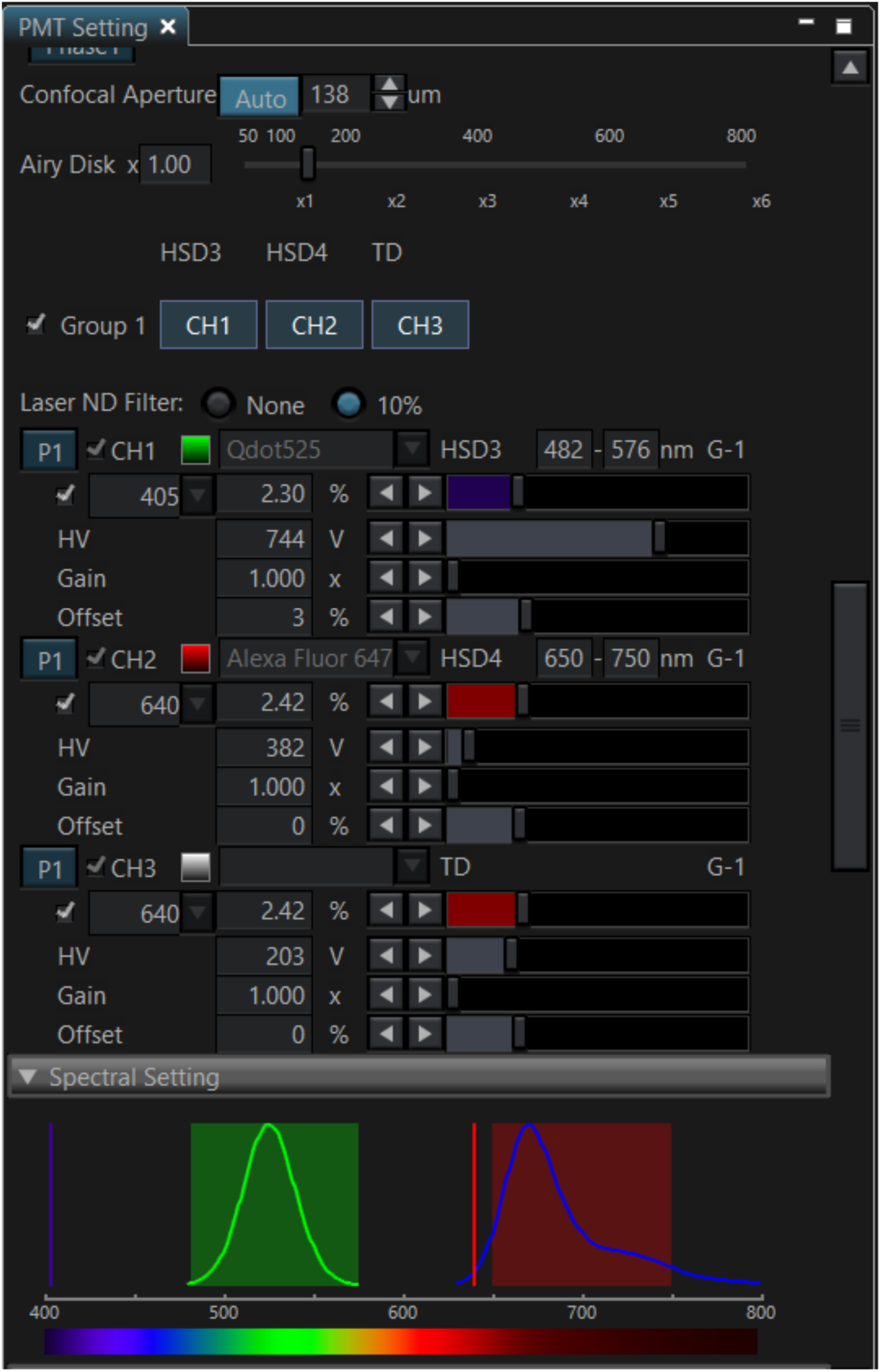
Sample instrument settings for Olympus FV3000 confocal microscopy studies. Rosindol signal (IND4) was detected via 405 nm excitation laser. WGA-A647 membrane stain was detected via 640 nm excitation laser.

**Figure S29.**
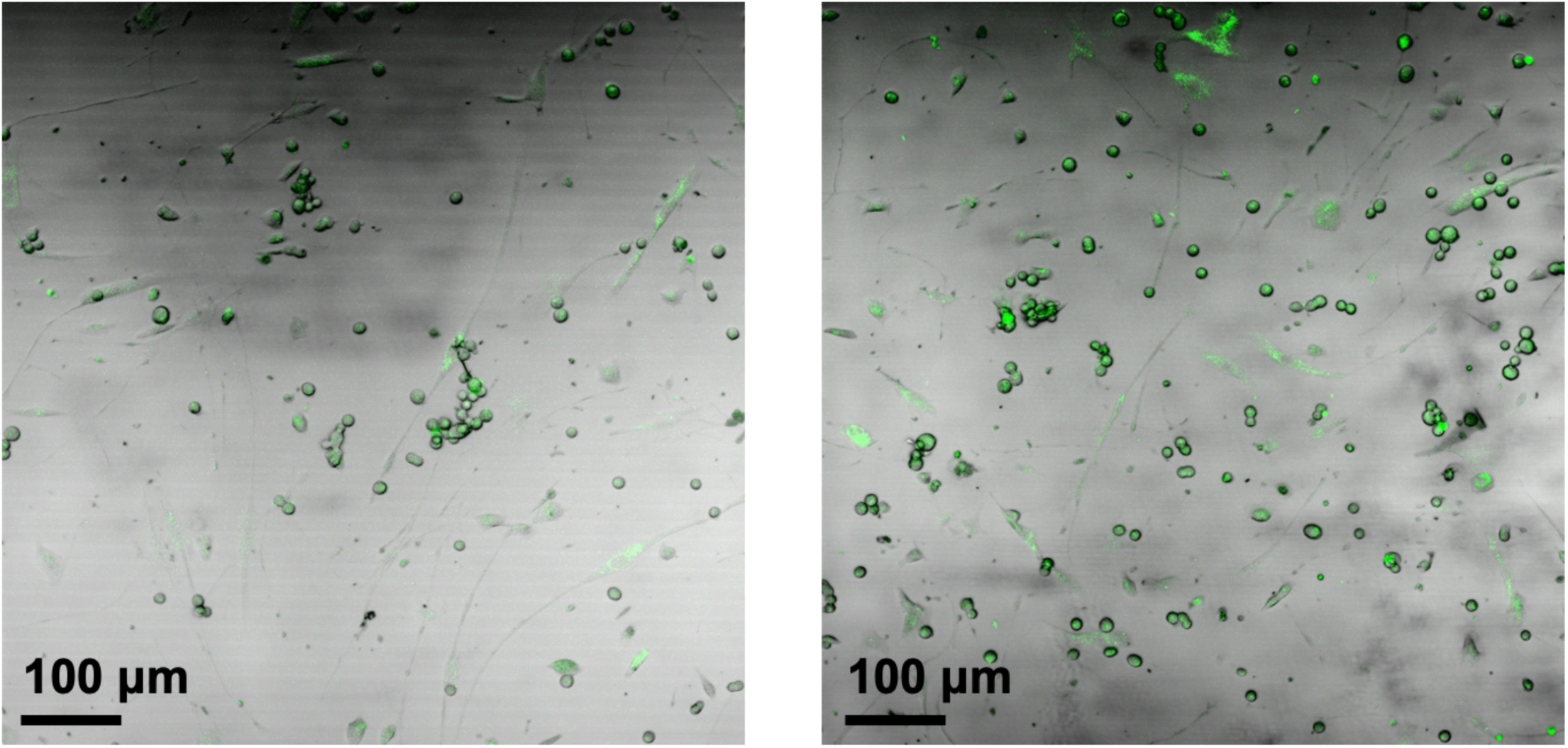
Confocal microscopy of a co-culture of Panc1/human dermal fibroblast cells treated with 100 μM Rosindol for 1 hour. Green signal represents IND4 as detected via 405 nm excitation laser. Panc1 cells appear spherical while fibroblasts possess elongated spindle-like morphology.

**Figure S30.**
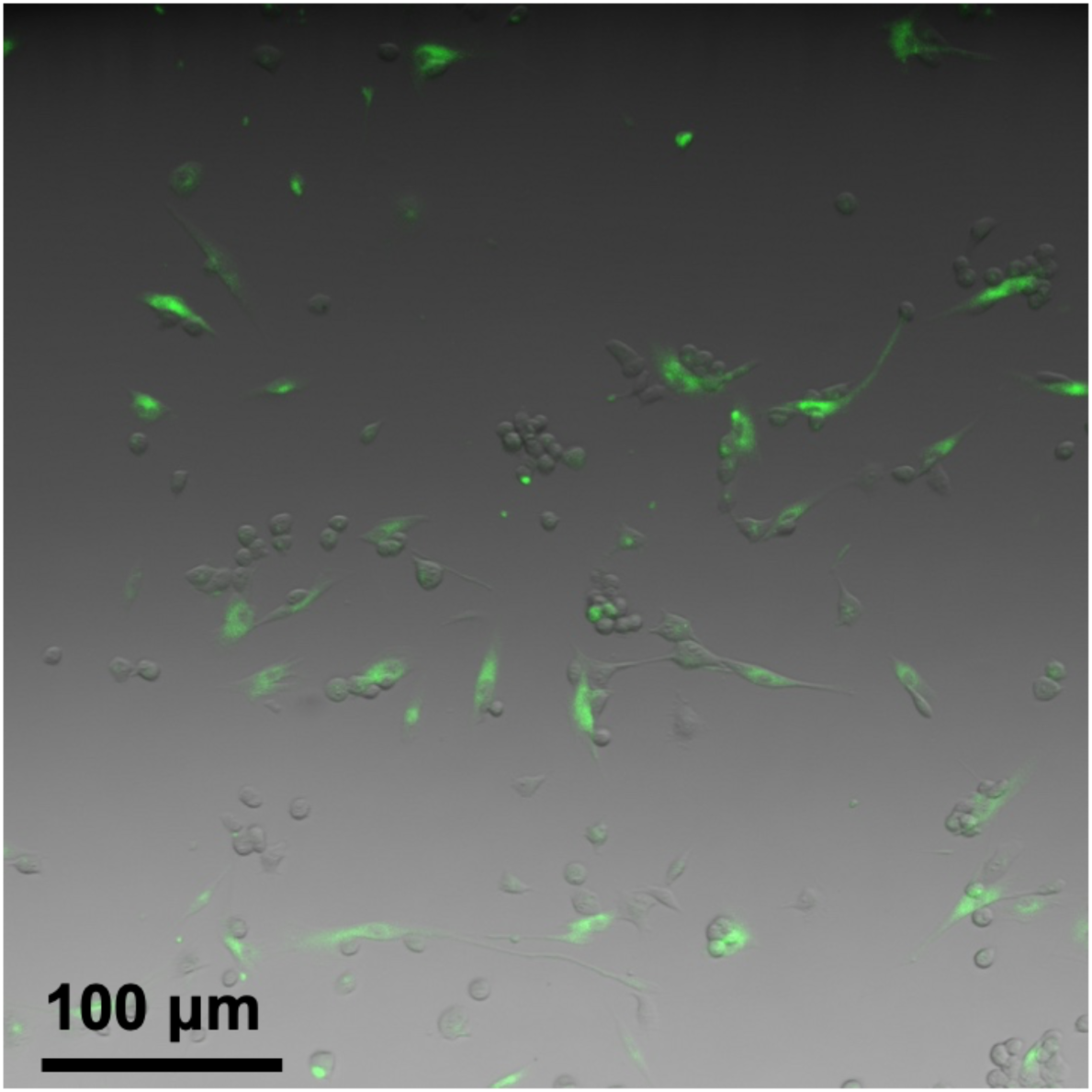
Confocal microscopy of a co-culture of Panc1/human dermal fibroblast cells treated with 10 μM H2D for 1 hour. Green signal represents DCF as detected via 488 nm excitation laser. Panc1 cells appear spherical while fibroblasts possess elongated spindle-like morphology.

**Figure S31.**
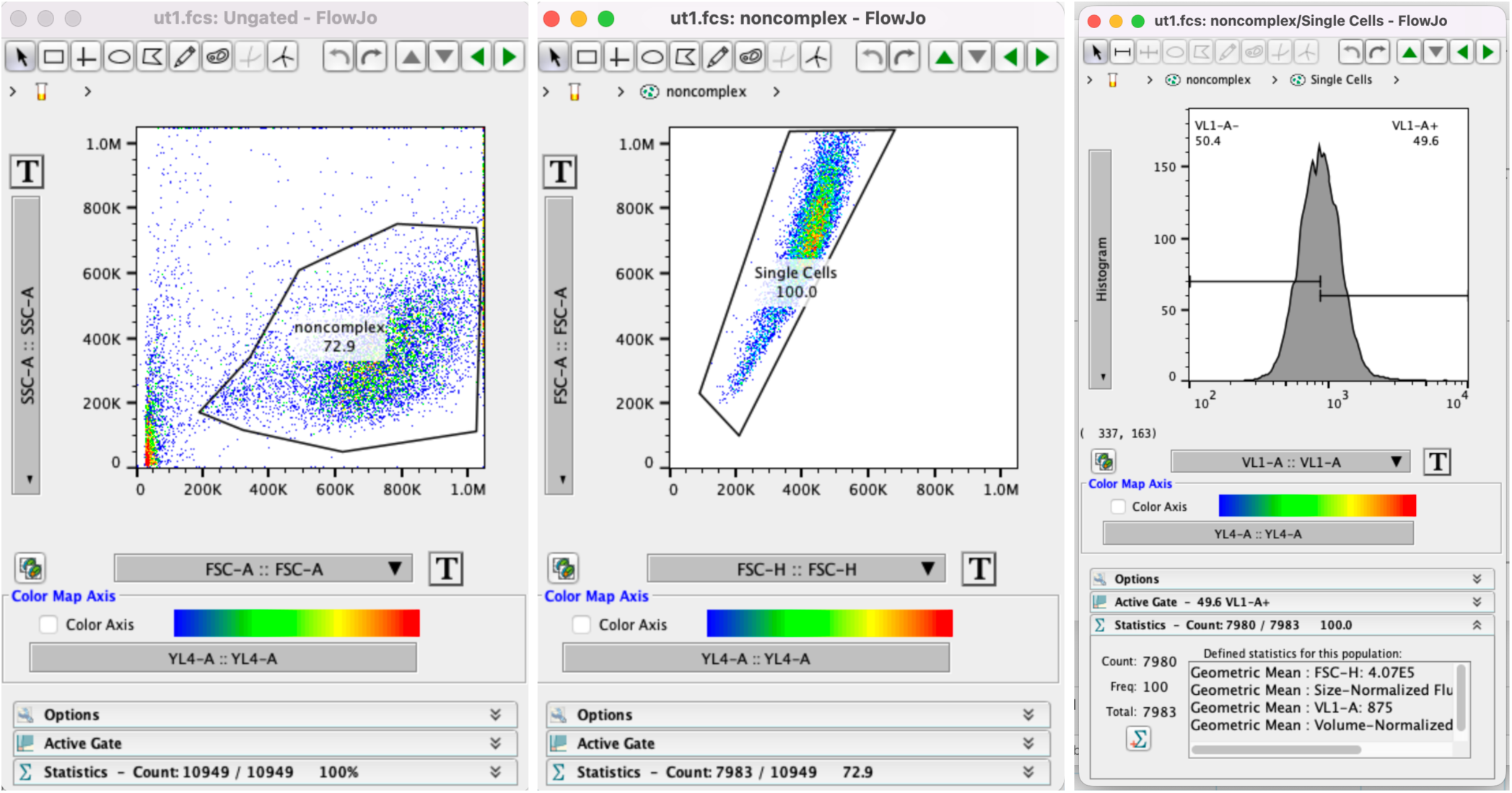
Sample forward and side scatter gating parameters for pancreatic cells analyzed via flow cytometry. Rosindol signal (IND4) was detected via 405 nm excitation using the laser VL-1.

**Figure S32.**
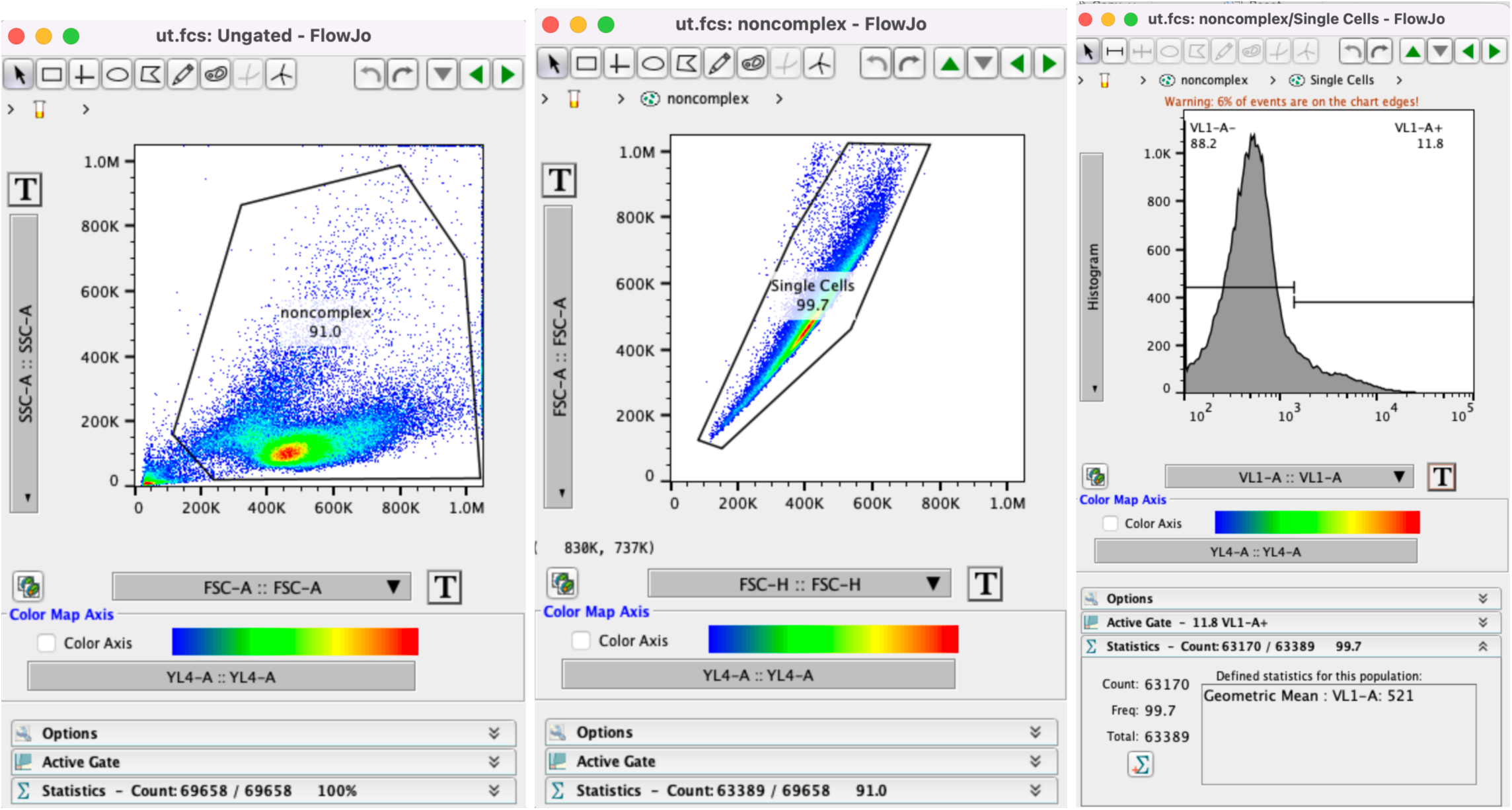
Sample forward and side scatter gating parameters for HL-60 neutrophils analyzed via flow cytometry. Rosindol signal (IND4) was detected via 405 nm excitation using the laser VL-1.

**Figure S33.**
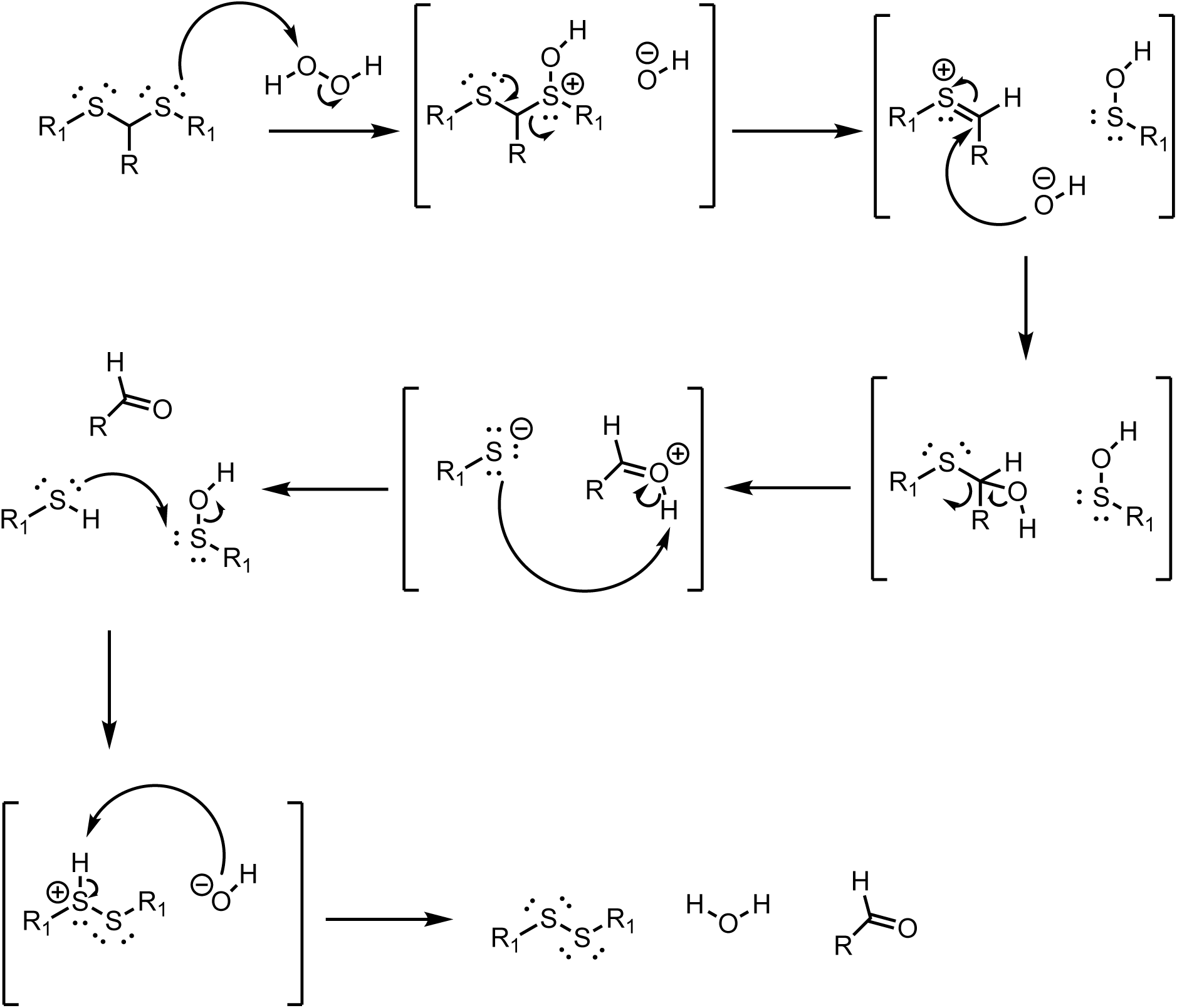
Mechanism of thioacetal oxidation via hydrogen peroxide. Adapted from Liu & Thayumanayan^51^.

**Figure S34.**
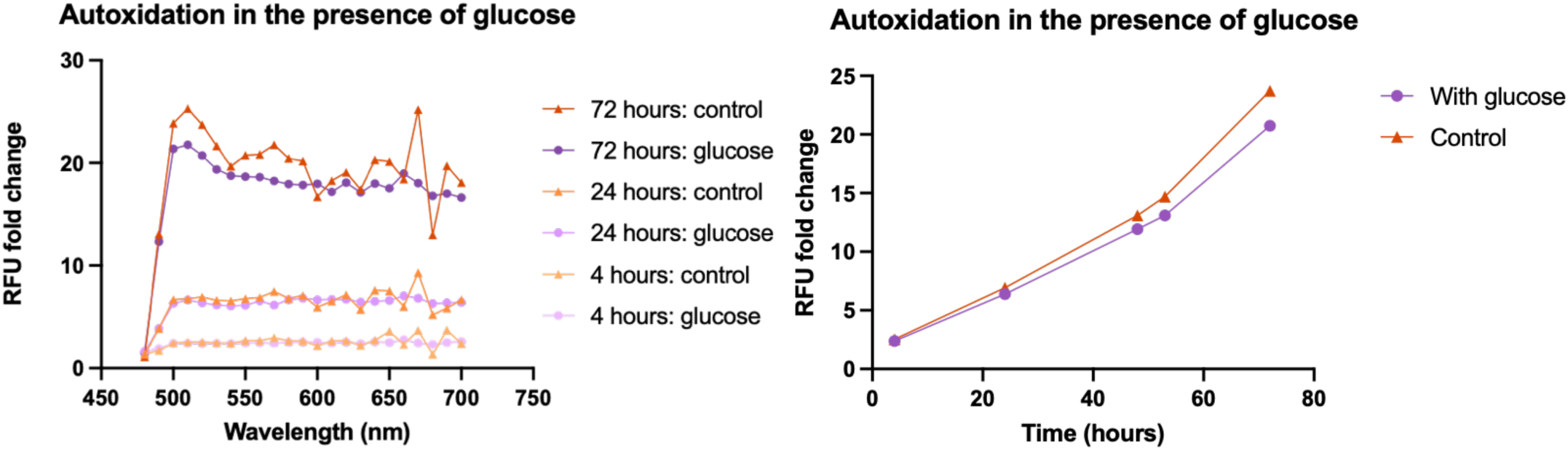
Oxidation of 10 μM boronate probe PF1 in the presence of 20 mM glucose. The data suggests that the presence of glucose alone does not cause fluorogenic oxidation of the boronate ester. Standard cell culture media (e.g. DMEM) glucose concentration is 25 mM.

**Figure S35.**
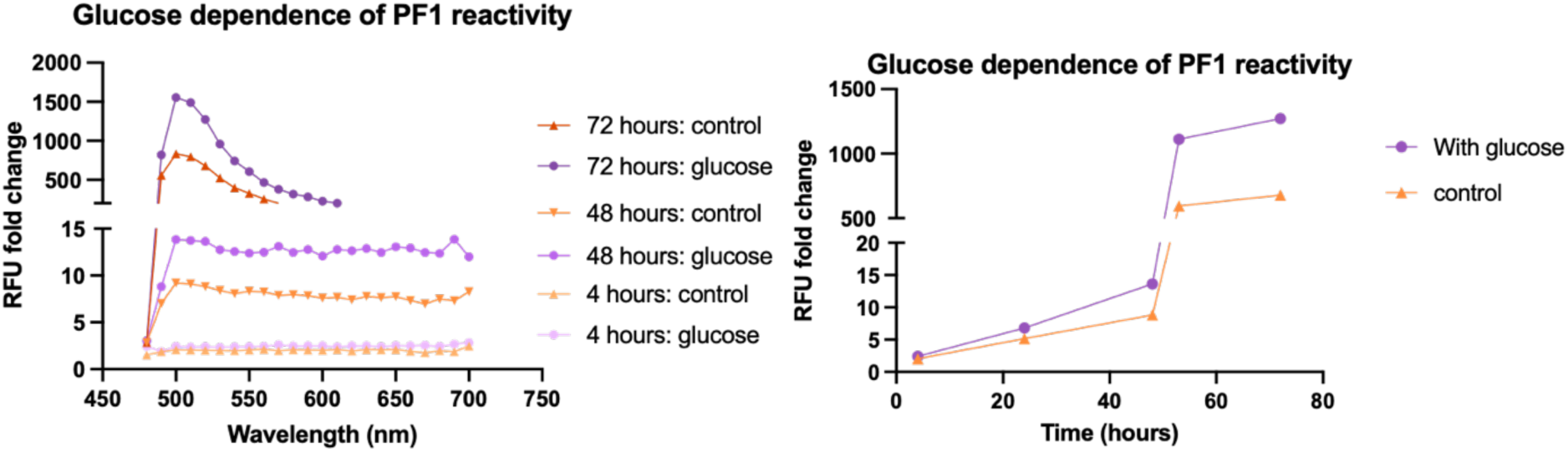
Oxidation of 10 μM boronate probe PF1 in the presence of 20 mM glucose and 100 μM H_2_O_2_. The data suggests that the glucose-bound probe is more reactive with H_2_O_2_, resulting in ROS signals dependent on glucose concentration. Standard cell culture media (e.g. DMEM) glucose concentration is 25 mM.

